# Combining Glycosidases and Nanopore Technology for Glycan Sequencing

**DOI:** 10.1101/2024.11.28.625867

**Authors:** Guangda Yao, Bingqing Xia, Fangyu Wei, Jiahong Wang, Yuting Yang, Shengzhou Ma, Wenjun Ke, Tiehai Li, Xi cheng, Liuqing Wen, Yi-tao Long, Zhaobing Gao

## Abstract

The efficient characterization of glycan structures remains a critical challenge. Direct glycan sequencing technologies with improved sensitivity and throughput are still needed. Here, we propose a glycan sequencing strategy based on glycosidase -assisted nanopore sensing. We used engineered nanopore α-hemolysin (M113R/T115A), which provided high-resolution discrimination of monosaccharide volume differences and sensitivity to variations in glycan chain length. By utilizing the specificity of glycosidases, we sequentially hydrolyzed the terminal residues of the glycan chains and detected the characteristic shifts in electrical signals generated by the translocation of hydrolysis products. The accuracy of recognition of hydrolysis fragments achieved over 90 % using machine learning. This allowed us to efficiently and conveniently determine the sequence of consecutive monosaccharide units in the glycan chains. Based on this principle, we conducted a proof-of-concept demonstration on actual samples and verified the accuracy via HPLC-MS. We achieved the sequencing of consecutive units in glycan chains, confirming the theoretical feasibility of glycosidase -assisted nanopore glycan sequencing. With future optimization, the development of arrayed nanopore glycan sequencing technology based on hydrolysis strategy will provide an innovative approach for rapid decoding of glycans, which could promote the progress of glycomics research and glycobiology.

## Introduction

Glycans play crucial roles in various biological processes such as cell differentiation, proliferation, immunity, aging, signal transduction, and migration.^1, 2^ They are composed of long chains of monosaccharides, with varying glycosidic linkages types, anomeric configurations, modification patterns, and branching structures.^3^ The complexity of these structures dictates their functions, and even minor changes in a glycan’s sequence can have significant effects.^4^ Therefore, an efficient and cost-effective strategy for glycan sequencing is essential for advancing glycoscience. Currently, nuclear magnetic resonance (NMR) spectroscopy and mass spectrometry (MS) are the primary techniques used for glycan structural analysis.^5^ Despite their widespread application and ability to provide some structural insights, these methods still face considerable challenges, including limited resolution, read length, accuracy, operational complexity, and time consumption.^6–8^ Compared to nucleic acid and protein sequencing, the development of glycan sequencing has notably lagged behind.^9^

Nanopore technology provides sequence information of analytes by detecting characteristic current signals generated by the translocation of target molecules through a nanopore,^10, 11^ which have been extremely successful applied in DNA sequencing.^10, 12^ Recently, efforts have been made to utilize nanopore technology for glycan recognition.^13^ Numerous attempts to improve the sensitivity and resolution of bio-nanopore detection have been reported, such as site-directed mutagenesis,^14, 15^ aptamer modifications,^16, 17^ and glycan tagging.^18^ Using these sensing strategies, detailed discrimination of minor differences such as building blocks,^16, 19, 20^ glycosidic linkages,^14, 17, 18^ length extension,^14, 18, 21^ branches,^18^ have been achieved. Nevertheless, these advancements demonstrate the potential of nanopores in glycan detection, yet they still fall short of deciphering the sequential information of continuous glycan units.^13^

In contrast to the linear linkage of 20 amino acids and 4 nucleotides, glycans comprise over 200 monosaccharide units, 5 linkage types, and multiple spatial conformations. The readout and analysis of the glycans-induced distinguishable signals are a tremendous challenge, even though machine learning.^18, 22^ Besides, unlike DNA and proteins with linear links and charges, glycan chains often have branches and low charge density and difficult to control during their translocation through nanopores.^23^ Therefore, addressing the issues of spatial extension and translocation is critical for achieving nanopore-based glycan sequencing. Here, we present an alternative strategy for glycan sequencing based on glycosyl hydrolase-assisted nanopore sensing. This approach involves hydrolyzing long glycan chains into smaller, rapidly analyzable monosaccharide units, offering high sensitivity, high resolution, and efficiency, though it has yet to be practically applied in nanopore glycan sequencing. Nanopore-based hydrolytic sequencing was initially proposed for nucleic acid sequencing,^24^ and recent studies have validated its feasibility for nanopore peptide hydrolytic sequencing with the aid of exopeptidases.^25 26^ The advantages of the hydrolysis sequencing strategy lie in effectively circumventing the issue of structural extension and its low requirement for control of translocation speed, only needing sufficient interaction between glycan fragments and the nanopore. Additionally, the hydrolysis process can be continuous, theoretically allowing unlimited read length. The vast array of naturally occurring glycosidases, with their high specificity for particular glycosidic bonds,^27^ offers an opportunity to obtain high-resolution structural information, thereby facilitating the development of nanopore-based glycan hydrolytic sequencing technology.

Building on our previous findings, we still utilized the α-HL(M113R/T115A) nanopore for acetylamino glycan sensing, which not only increased the frequency of molecular translocation but also provided higher resolution for distinguishing the volumetric differences of glycans.^14^ We conducted a proof-of-concept study for nanopore glycan hydrolytic sequencing using a heptasaccharide (Hepta) composed of LacNAc repeat units as a model glycan. As exoglycosidase were sequentially added, monosaccharide cleavage occurred at the reducing end of the glycan chain, resulting in distinct current signal shifts.^27^ Using two specific exoglycosidase, we preliminarily achieved sequential reading of five continuous building blocks and glycosidic bonds. This assay enables real-time detection of hydrolysis products during glycan degradation and allows for glycan sequence inference. Our study provides the first evidence that the combination of specific glycosidases and nanopore sensing technology is feasible for analyzing the monosaccharide composition and glycosidic linkage of glycans, offering strong support for further development of nanopore hydrolytic sequencing technology.

## Results

Our previous study demonstrated that α-HL(M113R) is a crucial glycan sensing site capable of detecting poly-LacNAc series glycans.^20^ By introducing the T115A mutation in the secondary sensing region, we expanded the detection window, enabling differentiation of Pentasaccharide (Penta) to Decasaccharide (Deca).^14^ Here, we continued using the M113R/T115A nanopore and poly-LacNAc glycans as model glycans to validate a nanopore glycan sequencing strategy assisted by enzymolysis (Method 1 and 2). To achieve high sensitivity and capture rate of random molecules, we conducted measurements in 3 M KCl buffer (3 M KCl, 10 mM citric acid, pH 5.0) with continuous voltage application, which was reported to increase the capture rate via improving electroosmotic flow (EOF). By convention, the electrode grounding chamber of the measurement device was defined as the Cis chamber, with the opposite chamber defined as the Trans chamber. Positive voltage was applied to the Trans side, with M113R/T115A proteins added to the Cis side (Figure 1A, Figure S1A, Method 3). The M113R/T115A formed a stable channel at +100 mV (Figure 1C), exhibiting a Gaussian-fitted open pore current value of 269.50 ± 3.23 pA (*I*_0_) and a conductivity (G) of 2458.21± 16.96 pS (Figure S1B-C). Upon establishing this stable single channel (Figure S1D), Heptasaccharide (Hepta, Figure 1B) was added to the *Cis* side at final concentrations of 25 μM, 50 μM, 100 μM, and 200 μM. As depicted in Figure 1C, Hepta induced reversible and short-lived current blockade events. We extracted event characteristics such as Δ*I*_1_/*I*_0_, Dwell time, and standard deviation (Std) with Pynanolab software (Figure 1D, Method 3 and 4). The capture rate calculated as signal frequency (events number/ t_total_) increased linearly with concentration from 25 μM to 200 Μm (Figure S2, Figure 1E). But both of Δ*I*_1_/*I*_0_ and Dwell time tend to remain unchanged as concentration increased (Figure S3), which indicated that Hepta could pass through the pore. Taking the solubility of Hepta into consideration, 100 μM is the optimal concentration for following nanopore detection. We further optimized the voltage driving force by increasing the voltage from +40 mV to +120 mV at a concentration of 100 μM Hepta (Figure S4). The results showed that the capture rate tended to increased linearly as the voltage raised from +40 mV to +120mV (Figure 1F). However, the pore became unstable beyond +100 mV (Figure S4), leading us to set the detection system voltage at +100 mV. The Δ*I*_1_/*I*_0_ and Dwell time of 100 μM Hepta at +100 mV showed single-peaked fitting curves with mean values of 0.3745 and 148.9 μs, respectively (Figure S3). Besides, the scatter plots of Dwell time versus Δ*I*_1_/*I*_0_ for Hepta clustered into single group (Figure S3). These characteristic nanopore events indicate Hepta could passes through the M113R/T115A nanopore predominantly in a single form.

**Figure 1.**
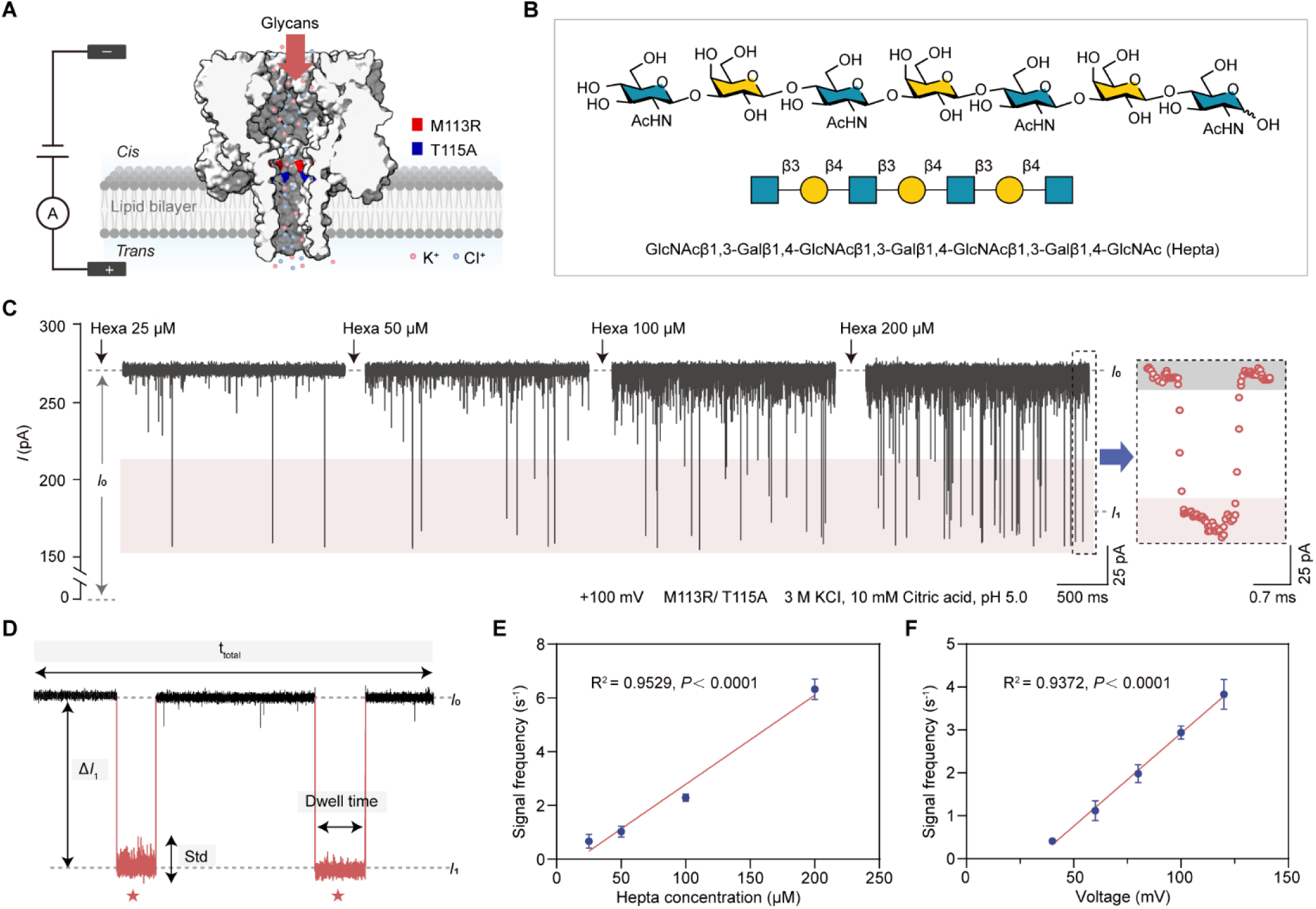
Nanopore-based glycan detection. (A) Schematics of the experimental setup with a α-hemolysin (M113R/T115A) inserted in lipid bilayer. The ionic current is recorded under a constant voltage at *Trans* side. (B) Chemical structure of Heptasaccharide (Hepta) and it illustration with Symbol Nomenclature for Glycans (SNFG). The blue square and yellow circle represent a N-Acetylglucosamine (GlcNAc) and a Galactose (Gal), respectively. (C) Detection of Hepta with gradient concentration. The ionic current traces recorded after addition of 25 μM, 50 μM, 100 μM, 200 μM Hepta show a tendency with increased current blockage events. The red star points denote an enlarged typical event (Right) in the left trace. (D) Depiction of event feature parameters used in glycan analysis. (E) Scatter plot of signal frequency versus concentration of Hepta, under applied +100 mV voltage at *Trans* side. (F) Scatter plot of signal frequency versus voltage applied at *Trans* side, the final concentration of Hepta is 100 μM. The formulas and adjusted R^2^ values and *P* values were computed on the basis of linear regression. All measurements were performed in a symmetric buffer (3 M KCl, 10 mM citric acid, pH 5.0), *n* ≥ 3 independent experiments.

We propose utilizing the specificity of glycosidases to release terminal units of glycan chains sequentially, enabling the stepwise decoding of individual monosaccharide units. Specifically, the first step involves cleaving the terminal building blocks (Monosaccharide, Mona) from the glycan chain. We can decode the glycan sequence information through the distinct electrical signal mediated by the released Mona passing through the nanopore, or the characteristic signal shift observed as the remaining glycan chain translocate through the nanopore. Thus, obtaining the reference fingerprints of the hydrolysis products and the corresponding exoglycosidases are crucial for the proof-of-concept of the hydrolysis strategy. According to theoretical predictions, Deca after the first hydrolysis step will yield a Nona and a Gal; Further hydrolysis of the Nona will produce an Octa and GlcNAc (Figure 2A). The remaining fragments will be Penta after 5 steps of enzymatic cleavage reaction. It could be convenient and efficient to distinguish hydrolysis products with single building block resolution through variation of one parameter such as Δ*I*_1_/*I*_0_ caused by the varying glycan length (Figure 2B). Under the current detection system, we first analyzed the standards of Deca hydrolysis products to establish fingerprints via M113R/T115A. Gal, GlcNAc, Penta, hexasaccharide (Hexa), Hepta (hepatsaccharide), octasaccharide (Octa), nonasaccharide (Nona), Deca were separately added to nanopore system, with final concentration of 100 μM. The results showed that the addition of Gal or GlcNAc did not cause any significant current blockade, even when the concentration increased to 1 mM (Figure S5). In contrast, the addition of 100 μM Penta, Hexa, Hepta, Octa, Nona, Deca produced reversible and sustained current blockade signals (Figure 2B). These observations suggested that the electrical signal displacement generated by the hydrolysis product of remaining glycan could be used to directly determine whether that hydrolysis has occurred at the reducing end of the glycan chain. By leveraging the specificity of exoglycosidase, we could accurately deduce the sequence of glycan chain from non-reducing end. Notably, the M113R/T115A nanopore does not respond to Mona, which helps avoid the interference with the signals of the hydrolyzed glycans. In detail, the scatter plots of Dwell time versus Δ*I*_1_/*I*_0_ for each glycan clustered into single groups (Figure 2C, Figure S6). The Gaussian-fitted values of Δ*I*_1_/*I*_0_ for Penta, Hexa, Hepta, Octa, Nona, Deca were 0.3607, 0.3421, 0.3745, 0.4684, 0.6350, 0.7378, with separation ratios of 0.2315, 0.4074, 1.6254, 3.2919 and 1.6618, respectively (Figure 2D, Method 4). The Exponential-fitted values of Dwell time of the six glycans were 235.9 μs, 143.7 μs, 148.9 μs, 139.0 μs, 97.12 μs, 108.0 μs, respectively (Figure 2E). The blockage amplitude is sufficient to distinguish these six glycans, and there was an approximate positive correlation between chain length and Δ*I*_1_/*I*_0_, indicating the translocation of the neutral glycan chain in the nanopore conforms to the size exclusion model. Overall, the distinct shifts in the electrical signal fingerprints of the three hydrolyzed glycan chains can be used to confirm glycan hydrolysis and infer the glycan structure.

**Figure 2.**
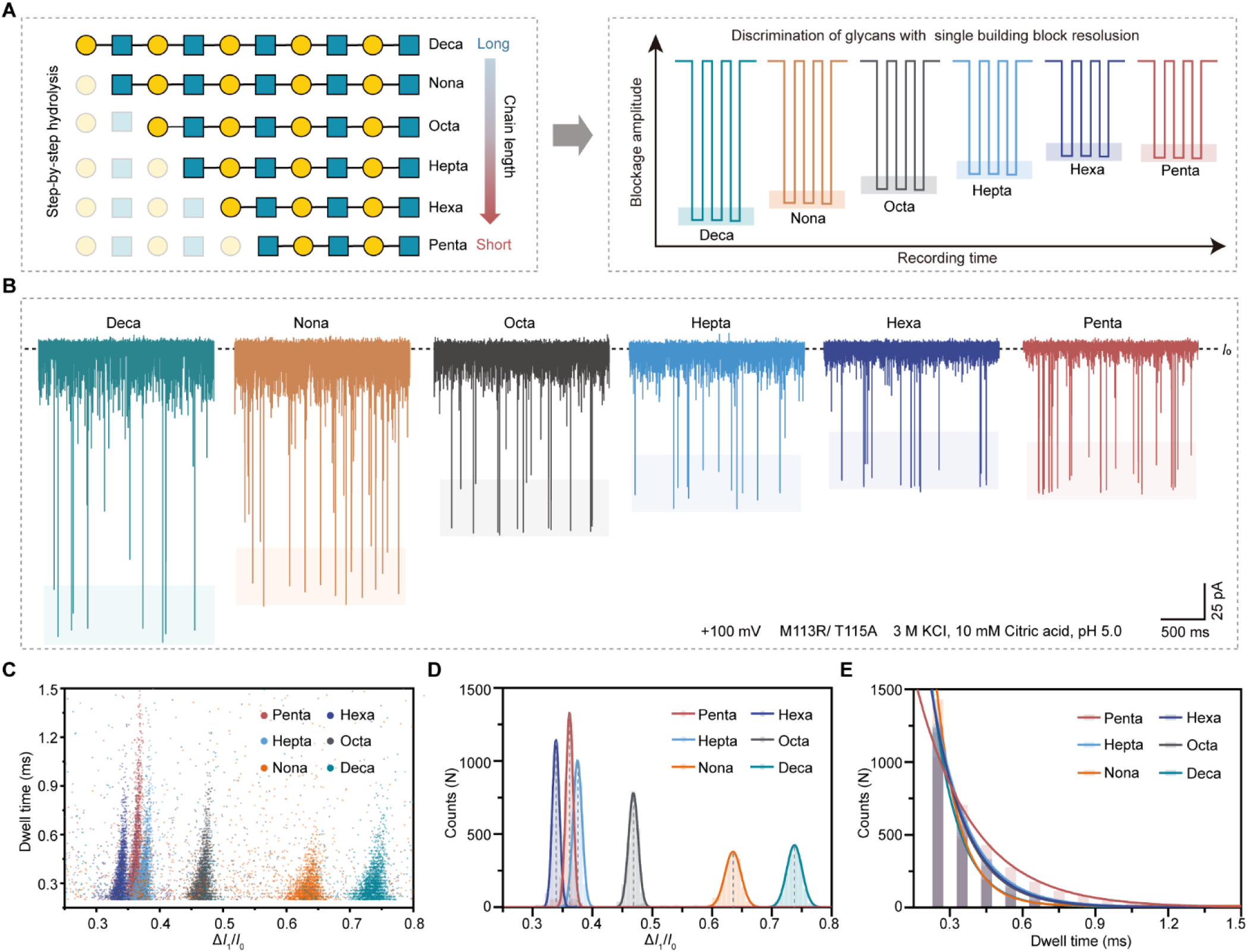
The fingerprints of glycans from Pentasaccharide (Penta) to decasaccharide (Deca). (A) Glycans from longer Deca to shorter Penta are depicted with Symbol Nomenclature for Glycans (SNFG). The blue square and yellow circle represent a N-Acetylglucosamine (GlcNAc) and a Galactose (Gal). The monosaccharides marked in diminished colors represent Gal and GlcNAc released by hydrolysis form the glycan non-reducing end. (B) Typical current traces of Gal and GlcNAc sensed by α-HL (M113R/T115A). (B) Schematic diagram of glycans discrimination with single building block resolution. (C) Typical ionic current traces of Deca, Nona, Octa, Hepta, Hexa, Penta sensed by α-HL (M113R/T115A). (C) The Dwell time versus Δ*I*_1_/*I*_0_ scatter plot of the six glycans (over 5000 events for each glycan). Δ*I*_1_/*I*_0_ was filtered ranging from 0.25 to 1.0, and the events with < 0.2 ms Dwell time were ignored. (D) Histogram of Δ*I*_1_/*I*_0_ distribution of Penta, Hexa, Hepta with Gaussian fitting. (E) Histogram of Dwell time distribution of the six glycans with exponential fitting. Each glycan was added to the *Cis* side with a final concentration of 100 μM. All measurements were performed in a symmetric buffer (3 M KCl, 10 mM citric acid, pH 5.0) under applied +100 mV at *trans* side. *n* ≥ 3 independent experiments.

Next, the model glycan Penta, composed of LacNAc repeating units, features Galactose (Gal) and N-Acetylglucosamine (GlcNAc) linked by β1,4-glycosidic bonds. The LacNAc repeating units are connected by β1,3-glycosidic bonds. Based on the structure, we purified two exoglycosidases specific for cleaving Galactose (GalH) and N-Acetylglucosamine (GlcNAcH) at the non-reducing end of poly-LacNAc glycans (Figure S7, Method 5 and 6). The enzyme activity and selectivity were assessed using thin-layer chromatography (TLC) with anisaldehyde staining. As shown in Figure 3A, Deca (S10) treated with GalNAcH for 10 minutes produced S9, which exhibited clear Nona bands with no detectable Octa bands. Following treatment of S9 with GlcNAcH for 10 minutes, S8 displayed clear Octa bands with no detectable Hepta bands. The TLC results indicated that after the first reaction, Deca was completely hydrolyzed to Nona, and after the second reaction, Nona was fully hydrolyzed to Octa. These results confirmed that the two exoglycosidases we obtained are specific and functional.

**Figure 3.**
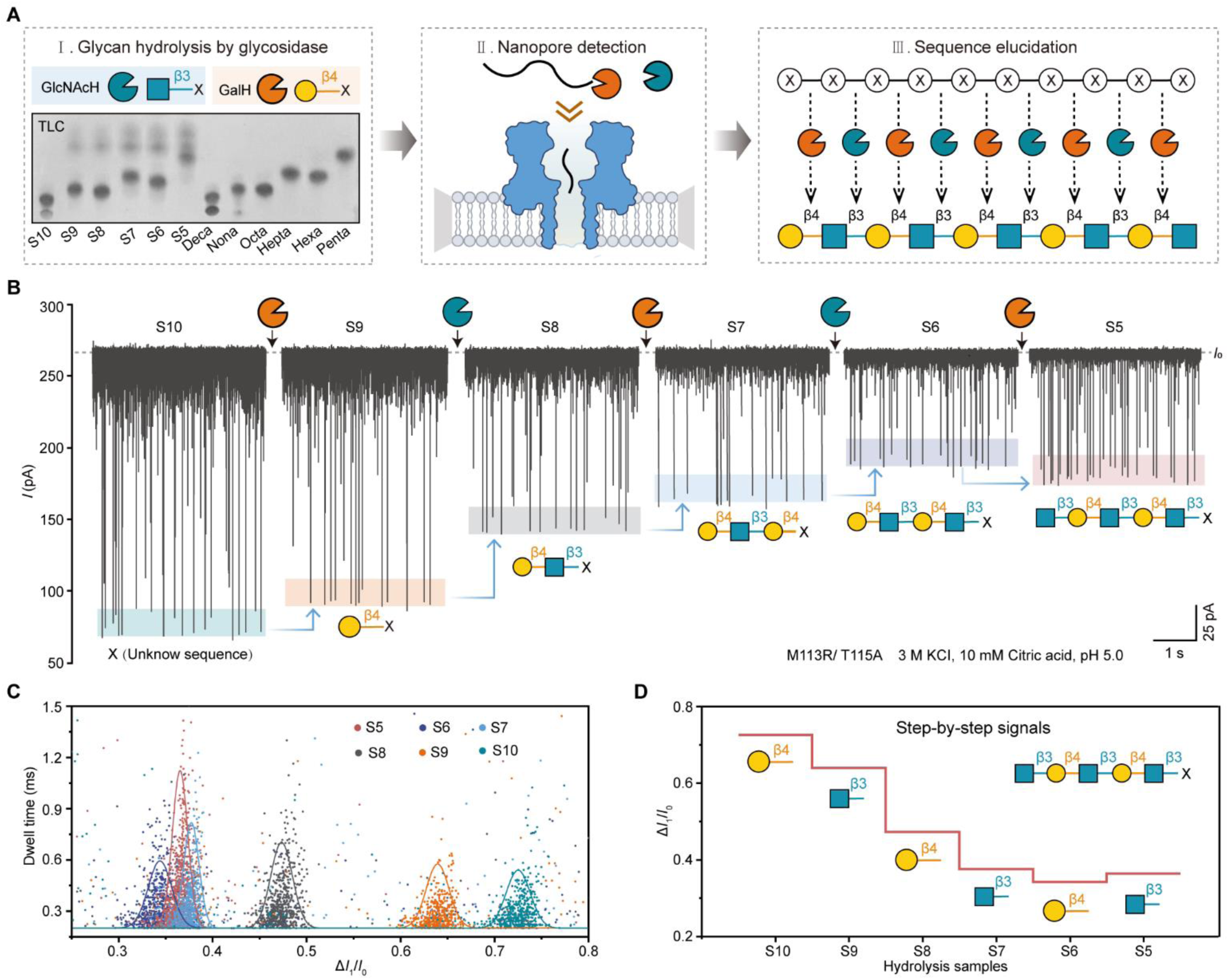
Nanopore-based glycan sequencing via terminal emzymolysis. (A) Three steps of nanopore glycan sequencing. Step 1 (Left), Stepwise hydrolysis of decasaccharide (Deca) using specific exoglycosidases of GlcNAcH (blue sector) and GalH (orange sector). The sample from S10 to S5 are the Deca and its stepwise hydrolysis products by GalH and GlcNAc. S10 is a solution of Deca (5 mM) mixed with inactivated 0.5 mg/mL GlcNAcH as negative control. The typical results of thin-layer chromatography (TLC) with anisaldehyde staining of S10, S9, S8, S7, S6, S5, compared with the standards solution of 5 mM decasaccharide (Deca), nonasaccharide (Nona), Octasaccharide (Octa), Heptasaccharide (Hepta), Hexasaccharide (Hexa), Pentasaccharide (Penta), (Developing agent, isopropanol: ammonia: water = 7:3:2). Step 2 (Middle), nanopore detection of samples from S10 to S5 after centrifugation. Step 3 (Right), glycan sequence elucidation based on the monosaccharide specificity and glycosidic bond preference of each exoglycosidase. X in circle represents the unknown monosaccharides. The arrows point to the cleavage sites on glycan nonreducing end. (B) Upper, representative ionic current traces of 2.6 μL S10, S9, S8, S7, S6, S5 sensed by α-HL(M113R/T115A). Bottom, stepwise glycan sequence elucidation based on the hydrolysis sample signals. (C) Dwell time versus Δ*I*_1_/*I*_0_ scatter plot the six samples form (B) (about 1000 events for each sample), with Gaussian fitting curve. (D) Horizontal step plot of Δ*I*_1_/*I*_0_ mean values of samples from S10 to S5. Each sample was added to the *Cis* side. All measurements were performed in a symmetric buffer (3 M KCl, 10 mM citric acid, pH 5.0) under applied +100 mV voltage applied at the *trans* side. *n* ≥ 3 independent experiments.

As an ultimate goal, we present a proof-of-concept study demonstrating glycan sequencing using strategies developed in this work. Mimicking real-world detection scenarios, the sequencing process was divided into three steps (Figure 3A). First, glycan hydrolysis via glycosidases (described in the Method 6 and illustrated in Figure S7). In brief, a solution of Deca (10 mM) were mixed with inactivated GalH as negative control and named as Sample 10 (S10). Another Deca solution were incubated with GalH at 37℃. After 10 min, the reaction was terminated via heating at 100℃ and collected as Sample 9 (S9). Then, the second exoglycosidase, GlcNAc, were added to S9, and the solution were collected as Sample 8 (S8) after 10 min incubation. S8, S7, S6 or S5 was prepared by GlcNAcH or GalH via similar procedure. All the samples were ultracentrifuged, and the supernatants were collected to be detected under the same conditions as glycan standards. Reversible current blockade events were recorded for at least 10 min continuously after addition of each sample (Figure 3B, Figure S8). Finally, glycan sequence elucidation. We aimed to obtain the sequence information based on the specificity of exoglycosidase. The events characteristic parameters of each sample were extracted to generate Dwell time versus Δ*I*_1_/*I*_0_ 2D scatter plots, in which the respective distinct clustering and significant relative movement between S10, S9, S8, S7, S6, S5 were observed (Figure 3C). Δ*I*_1_/*I*_0_ of samples from S10 to S5 were centered at 0.7262, 0.6400, 0.4728, 0.3761, 0.3425, 0.3640, with separation ratios of 1.3840, 2.8910, 2.0241, 0.6665, 0.4258 (Figure 3C-D). According to the step-by-step Δ*I*_1_/*I*_0_ variation (Figure 3D), and the specificity of these two enzymes and glycosidic bond preference, it is easy to inferred that the first monosaccharide at the non-reducing end of the substrate glycan is Gal, connected by a β1,4 glycosidic bond, and the second monosaccharide is Gal, connected by a β1,3 glycosidic bond. The five-step results illustrated that the model glycan chain is composed of β1,3-linked LacNAc units, which is consistent with the actual structure of Deca. To validate the accuracy of our nanopore detection system for hydrolysis products, we added standards from Deca to Penta with final concentration of 100 μM to hydrolysis samples from S10 to S5, respectively (Figure S9A). The events collected after addition of Penta, Hexa, and Hepta clustered within the scatter plot regions of corresponding samples (Figure S9C), with Δ*I*_1_/*I*_0_ centered at t 0.3630, 0.3750, 0.4703, 0.6357, and 0.7360, (Figure S9), highly consistent with pre-addition values (Figure S8). This further confirmed that the hydrolysis products were as expected, without interfering by-products. These results demonstrated the reliability of our nanopore-based glycan enzymolysis sequencing system. It is important to note, for unknown glycans, obtaining exoglycosidases and understanding their specificity is essential for hydrolysis assisted glycan sequencing. Theoretically, there are thousands of possible combinations of monosaccharides and glycosidic bonds. In practice, for specific classes of glycan, such as mammalian glycans, the types of monosaccharides and glycosidic bonds are limited. Therefore, it is feasible to develop a systematic kit of glycosidases for the corresponding glycan sequencing, especially with the assistance of high-throughput detection based on nanopore arrays.

Although the events generated by Deca, Nona, Octa, and Hepta during M113R/T115A detection are completely visually identifiable according to the significant differences in Δ*I*_1_/*I*_0_ Gaussian distributions. The lower separation ratio between the Gaussian distributions of events Δ*I*_1_/*I*_0_ for Hepta, Hexa, and Penta were 0.2315, 0.4074, which could lead to ambiguous discrimination of hydrolysis products based solely on Δ*I*_1_/*I*_0_ and reduce the accuracy of glycan sequencing. Machine learning can learn from the data through computerized algorithms, which has been widely used in previous reports for accurate identification of glycans with subtle differences by nanopores.^18, 19^ Moreover, with the assistance of machine learning algorithms, automatic identification of hydrolysis products could be achieved, which is beneficial for achieving rapid real-time glycan sequencing. In this study, in addition to Δ*I*_1_/*I*_0_, Dwell time express certain variation between different hydrolysis products, forming the base of machine learning. The entire machine learning workflow includes raw data acquisition, event feature extraction, and model training (Figure 4A). The event features generated by six glycan standards under the same nanopore detection conditions were extracted from the current-time trace, including Δ*I*_1_/*I*_0_ and Dwell time. Events with a time less than 0.2 ms were ignored to avoid high overlap. The Agglomerative Clustering algorithm was employed to filter for clustering events (Figure S10). A minimum of 1500 events were used to form the database for each glycan, with 80% used in the training set for model training and 20% used as the test set for model testing. Seven common machine learning models were evaluated including Support Vector Machine (SVM), K-NearestNeighbor (KNN), Decision tree, Adaboost, Neural Net, Naï ve Bayes, Random forest, with default parameters. To our satisfaction, Support Vector Machine (SVM), K-NearestNeighbor (KNN), Decision tree, Neural Net, Random forest achieving AUC value of > 0.95 and relative high accuracy of > 90% on the test set. Neural Net performed best by reporting the highest AUC value of 0.985 and highest accuracy of 90.9%, which was further used as model for automatic identification of hydrolysis sample. The feature of Δ*I*_1_/*I*_0_ contributed the most (93.43%) to the discrimination of six glycans, while Dwell time contributed 6.57% (Figure 3B). Then, the Neural Net model was applied on the testing set to generated confusion matrix, in which the accuracy of Penta, Nona, Hepta, Hexa, Penta were 86.7%, 98.0%, 88.5%, 100%, 100%, 100%, respectively, showing excellent classifying performance (Figure 3C). The learning curve was produced to assess the influence of size of training set on the accuracy (Figure 3D). Lastly, the Neural Net model was employed to identified the unlabeled events from the current trace of step-by-step hydrolysis sample detected by M113R/T115A. A minimum of 800 events of each sample after ignoration of 0.2 ms and filtration via Agglomerative clustering algorithm were fed to the model program. After prediction by this model, the signals from S10, S9, S8, S7, S6, S5 could be mainly attributed to Penta, Nona, Octa, Hepta, Hexa, Penta, respectively (Figure 4E-F). The assistance of machine learning improved the accuracy of hydrolysis product analysis, which could help to enhance the reliability of glycan hydrolysis sequencing based on nanopore. More importantly, automated data analysis and signal attribution provide feasibility for the realization of high-throughput glycan sequencing in the future.

**Figure 4.**
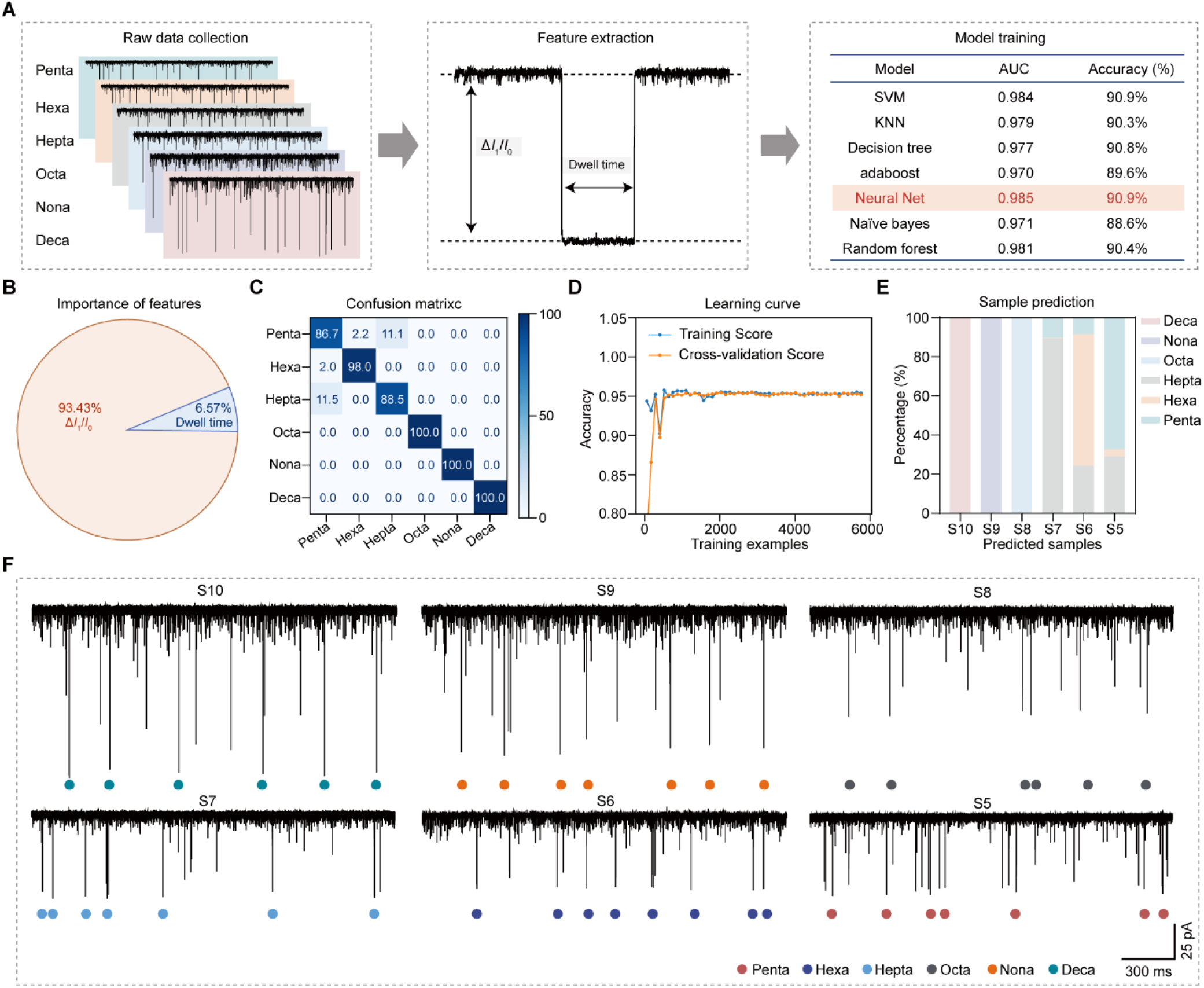
Prediction of glycan hydrolysis sample by machine learning. (A) The workflow of machine learning. A minimum of 1500 events of each glycan including Decasaccharide (Deca), Nonasaccharide (Nona), Octasaccharide (Octa), Heptasaccharide (Hepta), Hexasaccharide (Hexa), and Pentasaccharide (Penta) were selected to form a data set. Two features (Dwell time, Δ*I*_1_/*I*_0_) were extracted from blockage events to form a feature matrix. Seven models were separately employed and their corresponding AUC value and accuracies of test set were reported, among which the Neural set model (Marked in red) is the best performing model. (B) The importance of two features (Dwell time, Δ*I*_1_/*I*_0_) in model training. (C) The confusion matrix of six glycans classification generated by the Neural set model. (D) The learning curve of Neural set model produced by performing model training and validation using varying sample size. (E) Signal attribution of Deca hydrolysis sample (S10, S9, S8, S7, S6, S5) by the Neural set model, the signals marked with colored circle from S10, S9, S8, S7, S6, S5 were attributed to Penta, Nona, Octa, Hepta, Hexa, Penta, respectively.

Finally, to further validate the accuracy of our “hydrolysis-nanopore-reverse glycan structure” strategy, we confirmed the hydrolysis products using high-performance liquid chromatography (HPLC) and high-resolution mass spectrometry (MS). The HPLC column was eluted with acetonitrile/10 mM aqueous ammonium acetate pH 4.5 (60% acetonitrile) at a flow rate of 0.6 mL/min to analyze and distinguish the five glycan standards (Figure S11), with the retention time of 6.83±0.06 min, 7.73±0.06 min, 10.87±0.12 min, 13.13±0.23 min, 14.30±0.35 min, 17.67±0.47 min, 19.23±0.58 min, 24.03±0.58 min (Table S1). Then, the hydrolysis samples S10, S9, S8, S7, S6, and S5 were detected with the same HPLC method (Figure 5A, Table S2). The retention time of solution S10 was 24.10±0.46 min, belonging to Deca. The solution S9 was shown to eluting at 19.10±0.35 min and 7.73±0.06 min, for GalH removing the terminal Gal residue from Deca to obtain Nona. The hydrolysate S8 was shown to eluting at 17.57±0.29 min, 7.73±0.06 min and 6.82±0.03 min. Because GlcNAcH removing the terminal GlcNAc residue from Nona to obtain Octa. In this way, Hepta, Hexa, Penta were identified in S7, S6, S5, respectively. In addition, the mass spectrometry results demonstrated the generation of the expected product by using GlcNAcH and GalH (Figure 5B, Figure S12). The solution S10 was analyzed by ESI-HR and showed to Deca (*m/z* calcd for C_70_H_116_N_5_O_51_ [M-H]^-^ 1842.6632, found 1842.6632). The solution S9 was analyzed by ESI-HR and completely converted to Nona (*m/z* calcd for C_64_H_106_N_5_O_46_ [M-H]^-^ 1680.6121, found 1680.6121). ESI-HR profile of S8, S7, S6, S5 were showed to Octasaccaride (*m/z* calcd for C_56_H_95_N_4_O_41_ [M+H]^+^ 1479.5464, found 1479.5464), Hepta (*m/z* calcd for C_50_H_83_N_4_O_36_ [M-H]^-^ 1315.4792, found 1315.4792), Hexa (*m/z* calcd for C_42_H_70_N_3_O_31_ [M-H]^-^ 1112.3996, found 1112.3996), Penta (*m/z* calcd for C_36_H_60_N_3_O_26_ [M-H]^-^ 950.347, found 950.347) (Figure 5B, Figure S12). These results demonstrated that GlcNAcH and GalH glycosidases specifically performed stepwise cleavage of Hepta, validating the reliability of our nanopore hydrolysis sequencing strategy. Notably, M113R/T115A showed no response to Gal and GlcNAc, preventing interference from various hydrolysis products. Compared to TLC monitoring, our nanopore detection concentration was reduced by 100-fold. Additionally, our identification time was at least three times faster than MS and HPLC. Most importantly, the feasibility of designing a nanopore array allows the integration of the hydrolysis system with multiple units for current monitoring into a high-throughput nanopore sequencer. This integration offers the potential for rapid and efficient multistep hydrolysis sequencing.

**Figure 5.**
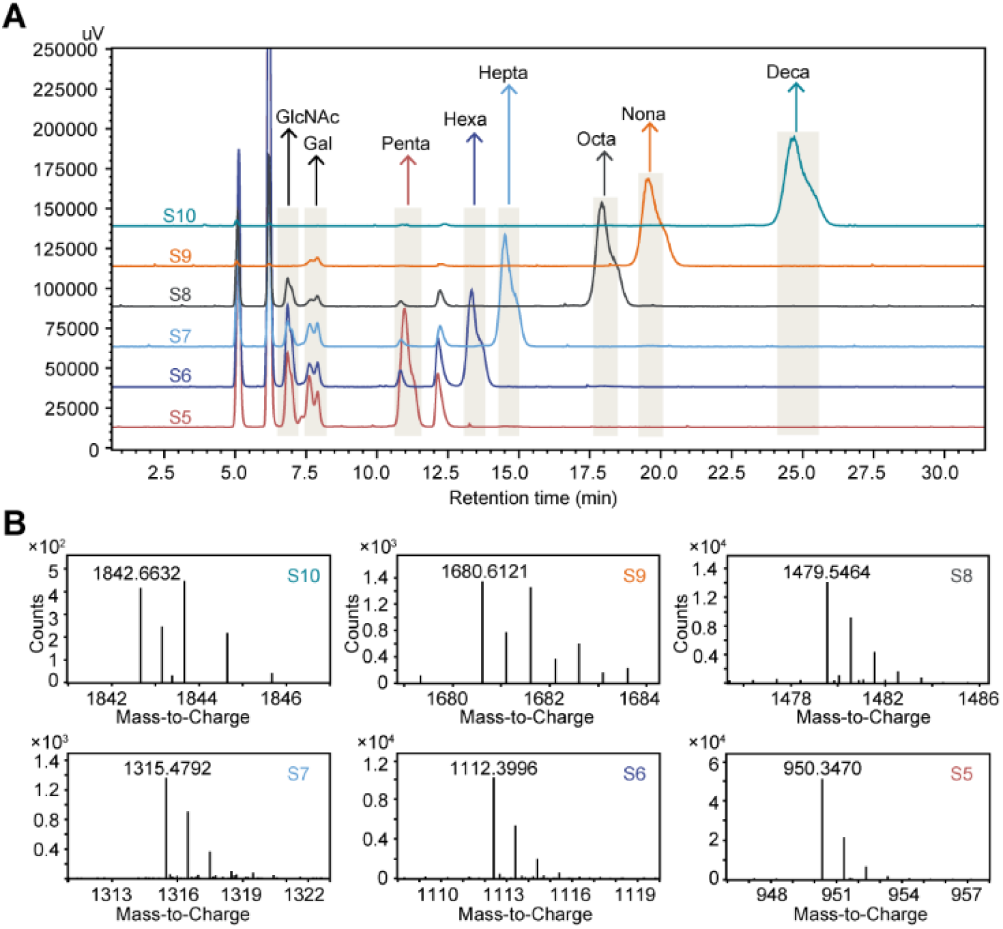
Verification of hydrolytic sequencing process. HPLC profile of the reaction solution samples, including sample 10 (S10), S9, S8, S7, S6, S5. The arrows marked in different color represent the glycans in samples identified based on the HPLC profile of the glycan standards. (B) ESI-HR profile of S10, S9, S8, S7, S6, S5, which were analyzed and showed to decasaccharide (Deca) (*m/z* calcd for C_70_H_116_N_5_O_51_ [M-H]^-^ 1842.6632, found 1842.6632), nonasaccharide (Nona) (*m/z* calcd for C_64_H_106_N_5_O_46_ [M-H]^-^ 1680.6121, found 1680.6121), octasaccharide (Octa) (*m/z* calcd for C_56_H_95_N_4_O_41_ [M+H]^+^ 1479.5464, found 1479.5464), heptasaccharide (Hepta) (*m/z* calcd for C_50_H_83_N_4_O_36_ [M-H]^-^ 1315.4792, found 1315.4792), hexasaccharide (Hexa) (*m/z* calcd for C_42_H_70_N_3_O_31_ [M-H]^-^ 1112.3996, found 1112.3996), Pentasaccharide (Penta) (*m/z* calcd for C_36_H_60_N_3_O_26_ [M-H]^-^ 950.347, found 950.347).

## Conclusion

In summary, we have demonstrated a strategy for sequencing glycans based on the glycosidase hydrolysis -assisted nanopore sensing. Our approach utilizes the specificity of glycosidases and nanopore sensing technology to monitor glycosidase-specific hydrolysis reactions, enabling rapid assessment by analyzing the monosaccharide composition and glycosidic linkage patterns of glycans. This strategy successfully interprets the sequential information of continuous monosaccharide units in glycan chains based on nanopore for the first time. In future work, specific glycosidases can be arrayed into a hydrolysis glycosidases system. By introducing unknown glycans into different enzymatic hydrolysis systems and real-time monitoring of any shifts in electrical signals before and after addition, we can directly determine which glycosidase is involved in the hydrolysis reaction. This allows us to reverse deduce the terminal monosaccharide module of the glycan chain. Repeating this process continuously will enable the determination of the entire glycan structure and de novo sequencing. The advantage of our system lies in its ability to directly judge hydrolysis by observing whether there is an electrical signal shift resulting from the hydrolyzed products.

This strategy utilizes glycosidase specificity to obtain information on the cleaved monosaccharide and linkage type, thus avoiding the need to establish fingerprint spectra for each monosaccharide and more complex glycans. While this study provides strong evidence for the feasibility of developing nanopore hydrolysis sequencing technology, there are some limitations. The system relies on the specificity of glycosidic bonds, but not all glycosidases exhibit such specificity, nor can all glycosidases be exhaustively covered. Enzyme engineering modifications in the future could offer a viable solution.^28^ Additionally, the relationship between glycosidase hydrolysis kinetics and characteristic bioelectrical signals, and the underlying mechanisms have not yet been elucidated, which would be beneficial for future qualitative and quantitative analyses and exploring the feasibility of de novo glycan hydrolysis sequencing. We anticipate that future developments of the platform will overcome the limitations of current sequencing procedures and provide an innovative technical approach for nanopore-based “glycan decoding,” promoting the discovery of glycan-based drugs and advancing glycoscience.^29–31^

## Materials

All of glycans in this research were purchased or synthesised by Wen-Lab. The chemical reagents are listed below:

### (1) Synthesis of glycans

Thin-layer chromatography (TLC) was performed on silica gel 60 F254 plates (Merck, MA) using *p*-anisaldehyde sugar stain. Molecular cloning reagents were from Invitrogen. Monosaccharides were purchased from BioChemSyn (www.biochemsyn.com, Shanghai, China). All other chemicals unless otherwise stated were purchased from Sigma or Carbosynth without further purification. ^1^H-NMR and ^13^C-NMR spectra were recorded on a Bruker 600-MHz NMR spectrometer (D_2_O as the solvent). High-resolution electrospray ionization (ESI) mass spectra were obtained using LC-MS (Thermo HPLC-Orbitrap Elite). Size Exclusion chromatography was performed using a column packed with Bio-Gel P-2 fine resins (Bio-Rad, Hercules, CA). HPLC analysis of compounds was performed on a Shimadzu LC-20A with a Waters XBridge BEH, Amide column, 5 µm, 4.6 x 250 mm. HPLC grade acetonitrile and water were purchased from Fischer.

### (2) Protein expression and purification

PageRuler™ Prestained Protein Ladder (Thermo Fisher, USA), isopropyl-β-D-1-thiogalactopyranoside (IPTG, Yeason, China), E.Z.N.A.® Plasmid DNA Mini Kit I (Omega Bio-Tek, China), Peroxidase AffiniPure Goat Anti-Mouse IgG(H+L) (Yeason, China), tryptone (Thermo Fisher, USA), agar (Sinopharm Chemical Reagent Co., Ltd, China),agarose (Yeason, China), Tris (Meilunbio, China), ampicillin sodium salt (Yeason, China), yeast extraction (Thermo Fisher, USA), ethanol (Sinopharm Chemical Reagent Co., Ltd, China), PrimeSTAR® Max DNA Polymerase (Takara Biomedical Technology (Beijing) Co., Ltd, China), HisSep Ni-NTA agarose resin 6FF (Yeason, China), dithiothreitol (DTT, Yeason, China), imidazole (Yeason, China), SDS−PAGE electrophoresis buffer powder (Yeason, China), HRP-conjugated Histag (Yeason, China), ethylene dinitrile tetra-acetic acid (EDTA, Sigma-Aldrich, USA). Super ECL Detection Reagent (Yeason, China), Coomassie brilliant blue G250 (Yeason, China)

### (3) Nanopore detection of glycans

Potassium chloride (Sigma-Aldrich, USA), citric acid (CA) (Sigma-Aldrich, USA), 1,2-diphytanoyl-sn-glycero-3-phosphocholine (DPhPC) (Avanti Polar Lipids, USA), chloroform (Sinopharm Chemical Reagent Co., Ltd, China), decane (Sigma-Aldrich, USA).

## Methods

### (1) The preparation of glycans by enzymatic synthesis

All glycans were enzymatically synthesized as reported previously.^14^ The structural information of glycans are described below:

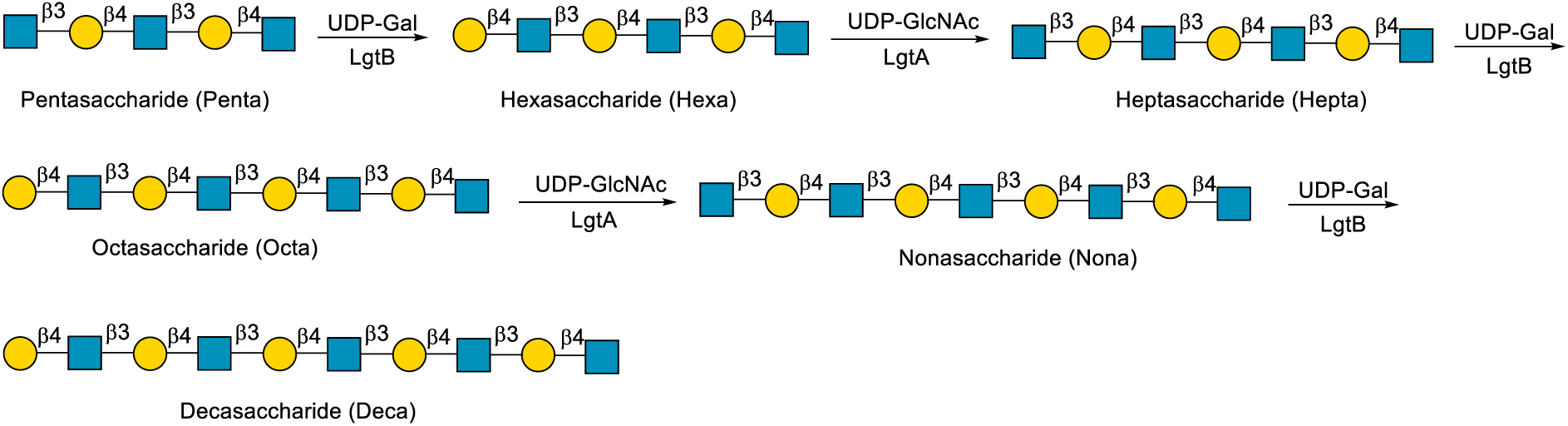

The glycans were prepared by enzymatic synthesis. The decasaccharide was synthesized from pentaccharide. A β14Gal was added by using a β-1,4-galactosyltransferase (LgtB) from *Neisseria meningitides* and a β13GlcNAc was added by using a β-1,3-N-acetylglucosaminyltransferase (LgtA) from *Helicobacter pylori*.

### (2) Nanopore protein preparation

Considering the reference sequence (NC_007795.1), the gene of α-hemolysin (α-HL, wild type) was synthesised by Beijing Genomics Institute (BGI, China) and inserted into the pEASY vector with a 6× His tag at the C-terminus for subsequent protein purification. The mutant plasmid (M113R/T115A) was constructed by polymerase chain reaction (PCR) using the high-fidelity PrimeSTAR® Max DNA polymerase with α-HL (wild type) -pEASY as the template under the following conditions: 98℃ for 15 seconds; 55℃ for 15 seconds; 72℃ for 210 seconds for 30 cycles. After first checking of PCR products by gel electrophoresis, they were treatede with 1 μl DpnⅠ (5 μl 10× QuickCut Green Buffer) at 37℃ for 15 min to digeste template. Then, the mutant products were transformed into *E.coli* trans5α competent cells (CD201, TransGen Biotech, China) by heat-shock and grown on LB plate, which containing the ampicillin (100 μg/mL) for resistance selection. Single colony was picked on the following day for DNA sequencing to vertify that the mutations had been constructed successfully. The vertified mutations could be used for the next nanopore protein purification via extracting plasmids with the Plasmid DNA Mini Kit (D6943-02, Plasmid Mini Kit I, USA).

The obtained α-HL (M113R/T115A) was transformed into *E.coli* BL21(DE3) pLysS competent cells (CD701-02, TransGen Biotech, China) and then grown on LB plate, from which single colony was picked for DNA sequenced (BGI, China) to confirm the sequence. The vertified colony was first amplified in 30 mL LB medium with 100 μg/mL ampicillin at 37℃ and 220 rpm for 14-16 hour. Subsequently, the cell suspension culture was transferred to 1.5 L LB with 100 μg/mL ampicillin medium for continued growth under 37℃/220 rpm. When the OD600 of the system reached 0.6-0.8, nanopore protein expression could be induced by 1 mM IPTG at 22 °C/220 rpm overnight. The cell suspension was centrifuged at 4℃ and 4000 rpm for 20 min to collect the pellet, and then the pellet was resuspended with TBS buffer (1 mM DTT) and lysed by high-pressure homogenizer for 3-5 min at 4℃. The soluble fraction obtained from cell fragments via centrifugation at 4 °C/13,000 rpm for 30 min was filtered with a 0.22 μM Millipore filter. The pre-equilibrated Ni-NTA resin (1 mL) with TBS buffer was mixed with the clarified supernatant and shaken for 1 h at 4℃ being transferred to the column. Linear concentration gradient of imidazole (10-30 mM) was utilised on the Ni-NTA affinity resin column to wash unbound non-specific proteins and washed with 300 mM imidazole to elute the target protein (α-HL). Samples from each step of the process were finally determined by SDS-PAGE to vertify the molecular weight of the target protein.

### (3) Nanopore detection and current recordings

Briefly, 1,2-diphytanoyl-sn-glycerol-3-phosphocholine (DPhPC, Avanti Polar Lipids) could be utilised to form lipid bilayer with a pore size of approximately 60 μM on the polytetrafluoroethylene film, which separated the two sides of the chamber (the electrode grounding chamber of the measurement device was defined as the *Cis* chamber, with the opposite chamber defined as the *Trans* chamber). Typically, 260 μL electrolyte solution (3 M NaCl, 10 mM citric acid, pH 5.0) was added to two sides, while α-HL (M113R/T115A) protein and the analytes were added to the *cis* side with a bias voltage of +100 mV applied to the *trans* side. When a stable open-pore current was observed, indicating that the nanopore protein α-HL (M113R/T115A) had been stably inserted into the lipid bilayer. The data were collected by nanopore instrument Cube-D2 (https://zenodo.org/records/11609574) after the addition of the analytes to the *cis* side, where the ionic current signals were low-pass filtered at 5 kHz and sampled at 50 kHz. The data were collected by nanopore instrument.

### (4) Data Analysis

All the current traces of single-molecule events were processed with PyNanolab software (https://doi.org/10.5281/zenodo.11383973). Current signals were analysed by GraphPad Prism 9 (Insightful Science, USA) and Origin 2022, and all data were presented as mean ± SEM. Characteristics of blockage events in current traces were extracted by PyNanoLab, including the blockage amplitude (ΔI), the event dwell time (τ) and Standard Deviation (Std) with ignore duration of 0.1 ms and ignore ΔI of 0.25. The data were analysed via GraphPad Prism 9 and Origin 2022 for blockage events feature fitting scatterplots, histograms and curve fitting. The separation (S) of different analytes in the scatterplot could be derived using the following formulas.

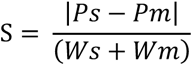

where Ps and Pm represent the peak values in the Gaussian fit of different analytes in the scatter plot, and Ws and Wm represent the half peaks widths in the Gaussian fit of different analytes.

### (5) Enzyme preparation

β-N-Acetylhexosaminidases from *Bacteroides fragilis* (GlcNAcH)^32^ and β1-4 Galactosidase from *Streptococcus pneumonia* (GalH)^33^ were synthesized artificially. Gene synthesis service was provided by Sangon Biotech (Shanghai, China). The gene was cloned into the pET-28a vector resulting in recombinant protein with six histidines at the protein N-terminal for further purification. The confirmed construct were subsequently transformed into E.coli BL21 (DE3) for protein expression. The His-tagged proteins were purified by using a Ni-NTA agarose column. Protein concentration was determined by the BCA Protein Assay Kit.

### (6) The hydrolysis test of pentasaccharide by using GlcNAcH and GalH

A 500 μL mixture containing 5 mM of substrate and 0.5 μg/μL GalH was heated in boiling water for 10 min. Insoluble impurities were removed by centrifugation (10000 rpm, 5 min). The supernatant was transferred to an eppendorf tube as Sample 10 (S10).

A 500 μL mixture containing 5 mM of decasaccharide and 0.5 μg/μL GaH was incubated at 37°C. When no starting material was observed on TLC, the reaction was quenched by heating in boiling water for 10 min. Insoluble impurities were removed by centrifugation (10000 rpm, 5 min). The supernatant was transferred to a tube as Sample 9 (S9). Afterwards, S8, S7, S6 or S5 was prepared by GlcNAcH or GalH via similar procedure.

### (7) General protocols for HILIC-HPLC analysis

The solution was analyzed by HPLC equipped with Essentia ELSD-16 detector using Waters XBridge BEH, Amide column, 5 µm, 4.6 x 250 mm. The column was eluted at 40°C with acetonitrile/10 mM aqueous ammonium acetate pH 4.5 (60% acetonitrile) at a flow rate of 0.6 mL/min, injection volume of 3 µL (5 mM).

## Supplementary Figures

**Figure S1.**
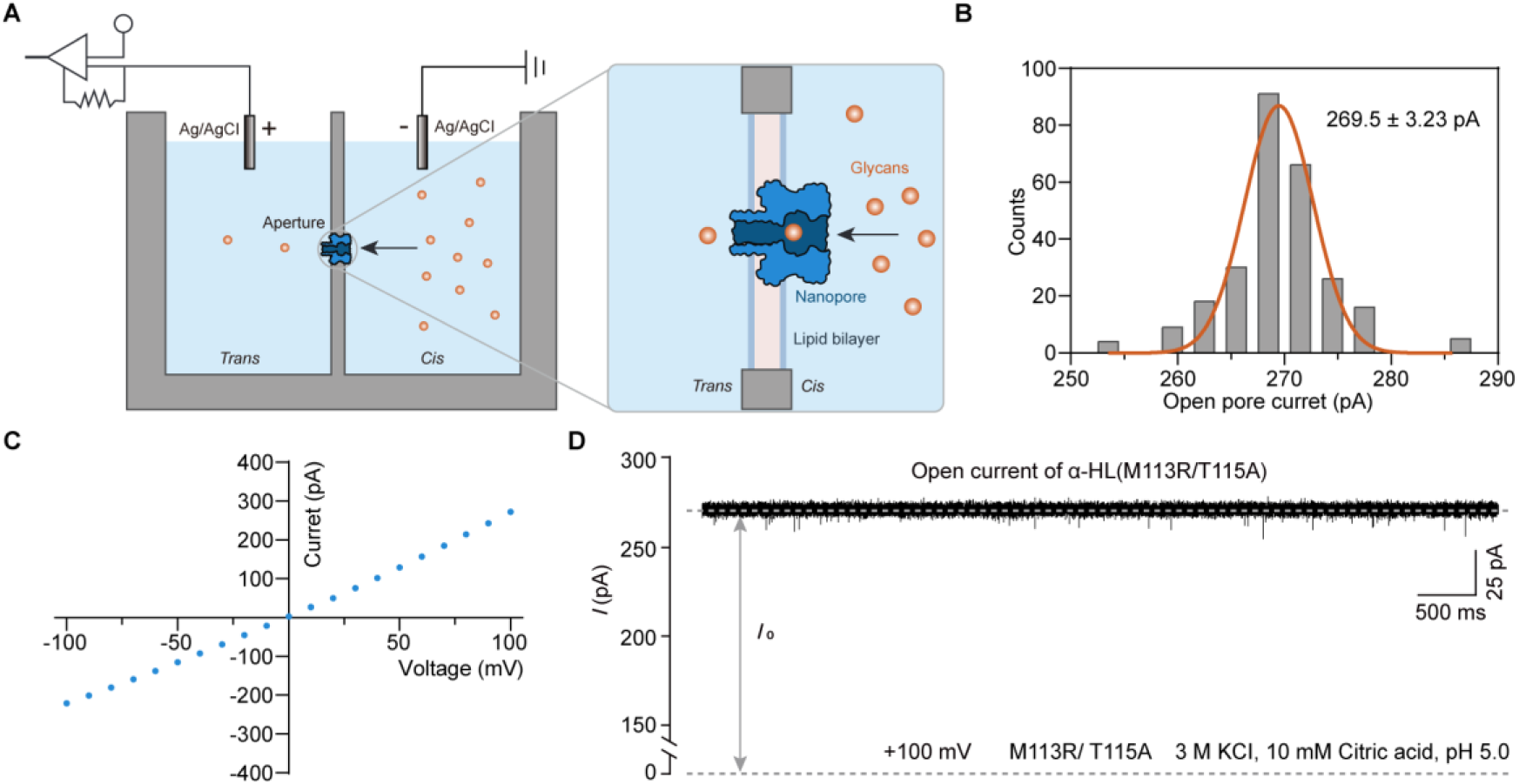
Nanopore-based glycan detection. (A) Schematic diagram of nanopore detection devices and enlarged view of the aperture part. The chamber was divided into two compartments and each containing 260 μL electrolyte solution (3 M KCI, 10 mM Citric acid, pH 5.0). The two compartments were connected through an aperture on which the lipid bilayer is formed. A constant voltage was applied at *trans* side. The glycan molecules translocate through the single nanopore form *cis* side to *trans* side. (B) Histogram with Gaussian fit of open pore current of M113R/T115A, +100 mV at *trans* side. (C)The current-voltage (*I*-*V*) curve of M113R/T115A (*n*≥3). (D) Representative ionic current trace of single M113R/T115A nanopore inserted in the lipid bilayer, under an applied voltage of +100 mV at tra*ns* side. All measurements were performed in a symmetric buffer (3 M KCI, 10 mM Citric acid, pH 5.0).

**Figure S2.**
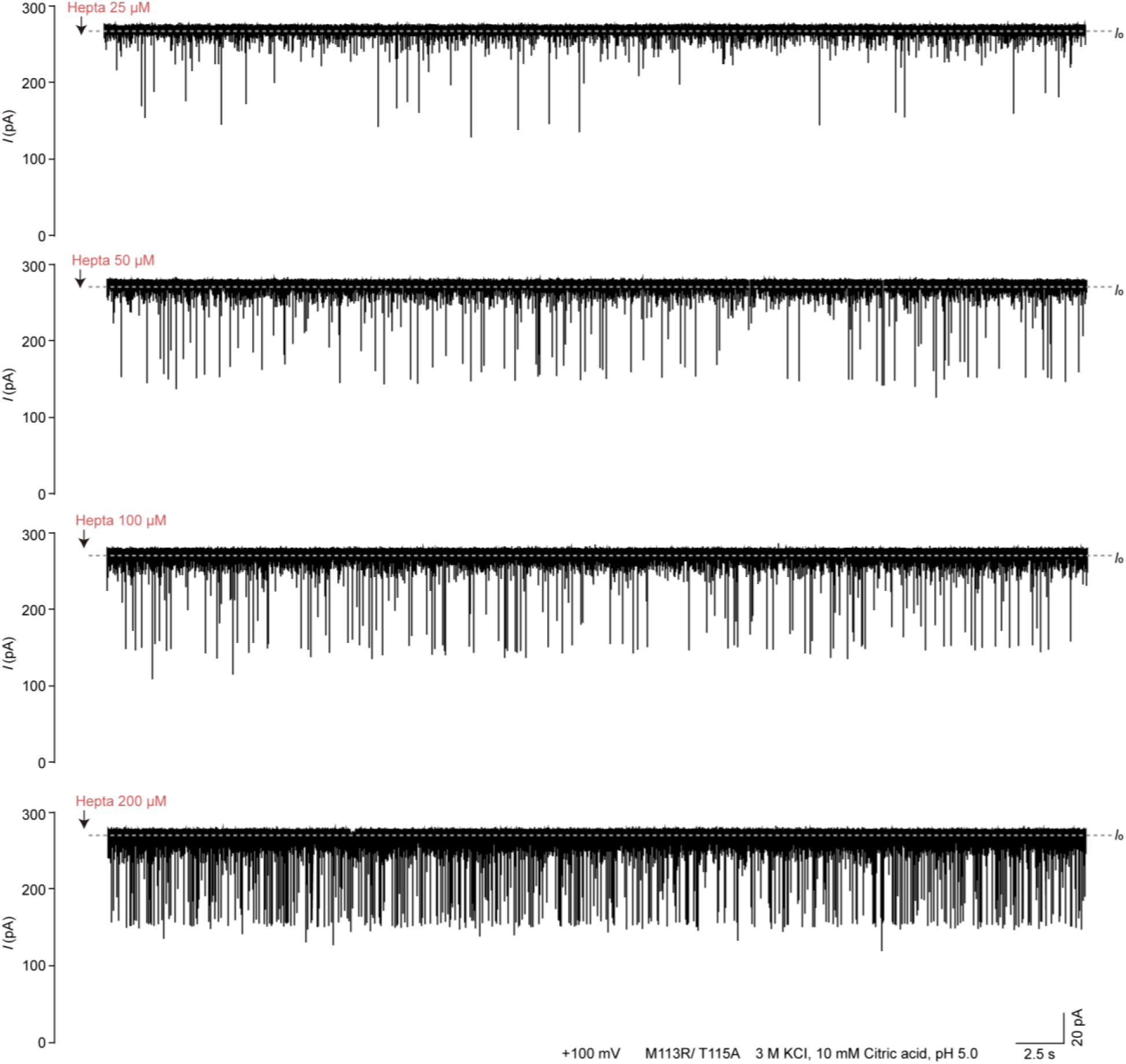
Detection of Heptasaccharide (Hepta) with gradient concentration. Representative ionic current traces after the addition of Hepta with gradient concentrations of 25 μM, 50 μM, 100 μM, 200 μM. *I*_0_ represents the open current of single M113R/T115A nanopore. The black arrows indicate the addition of glycan to the *cis* side. All measurements were performed in a symmetric buffer (3 M KCI, 10 mM Citric acid, pH 5.0), under an applied voltage of +100 mV at tra*ns* side. (*n*≥3)

**Figure S3.**
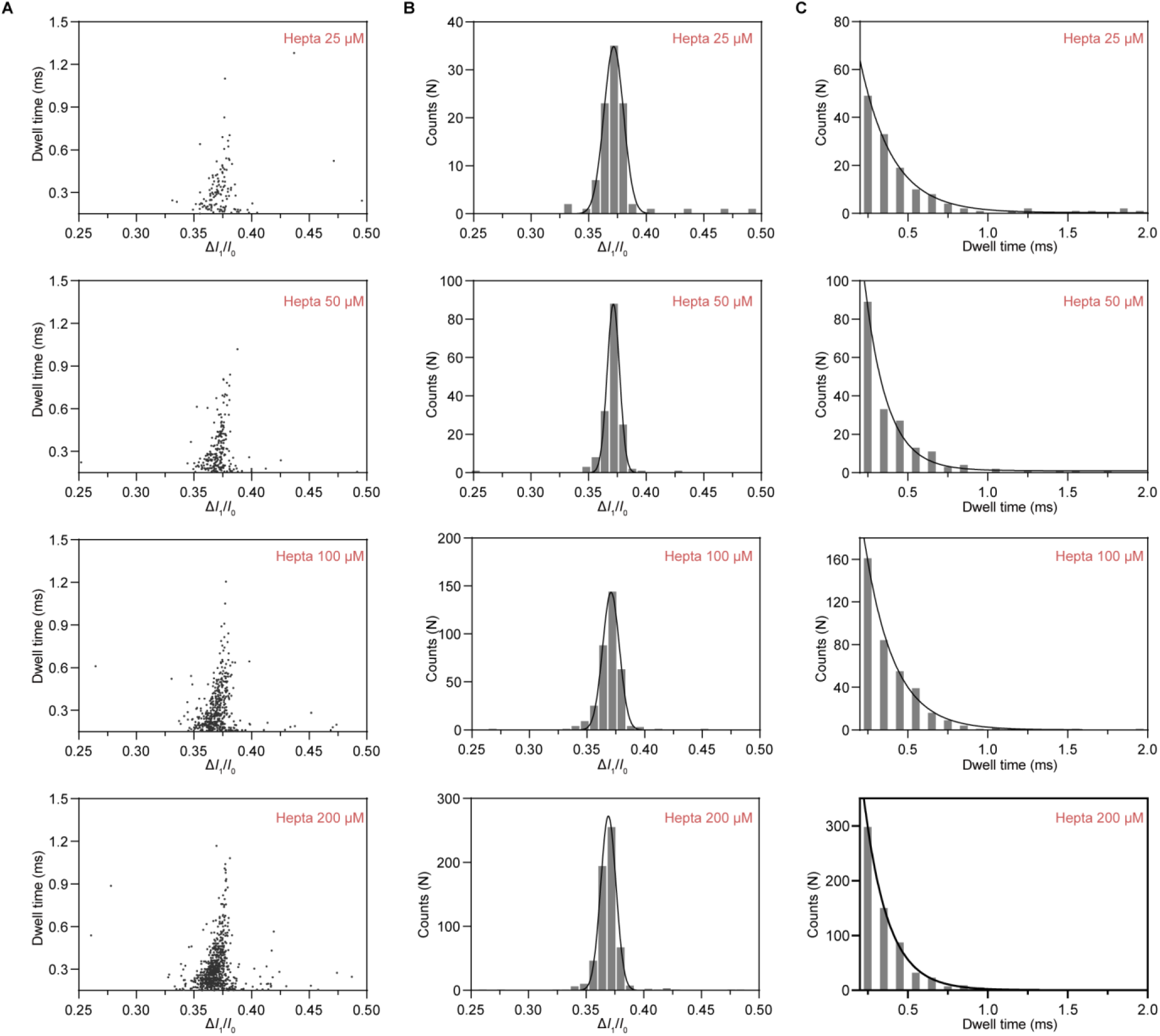
Statistics of events of Heptasaccharide (Hepta) with gradient concentration. (A) The scatter plots of Δ*I*_1_/*I*_0_ versus Dwell time of Hepta with concentration of 25 μM, 50 μM, 100 μM, 200 μM. (B) Histogram of Δ*I*_1_/*I*_0_ in (A) with Gaussian fitting. (C) Histogram of Dwell time in (A) with Exponential fitting. All measurements were performed in a symmetric buffer (3 M KCI, 10 mM Citric acid, pH 5.0), under an applied +100 mV voltage at tra*ns* side. (*n*≥3)

**Figure S4.**
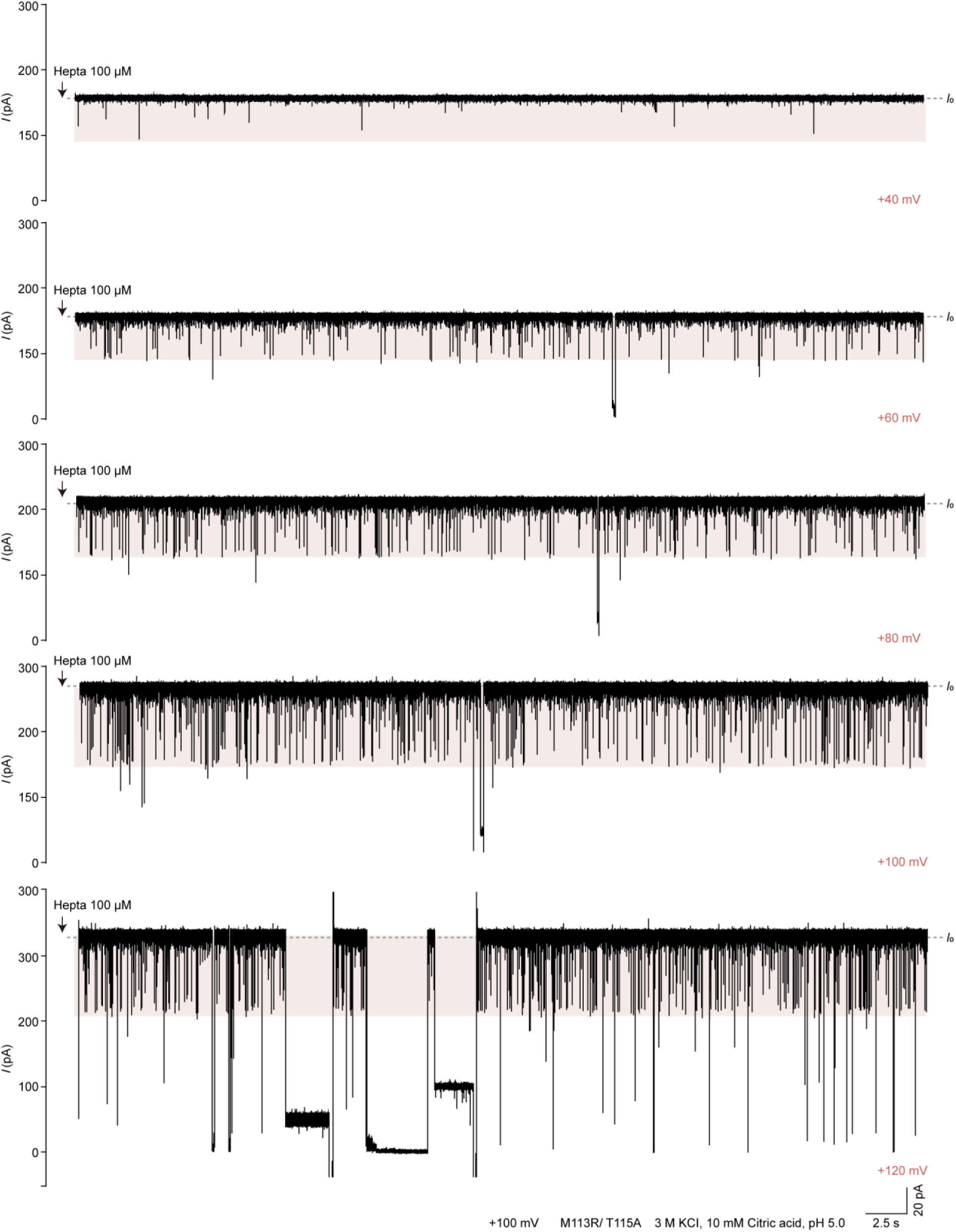
Detection of Heptasaccharide (Hepta) under gradient voltages. Representative ionic current traces after the addition of 100 μL Hepta to *cis* side, under gradient voltages of +40 mV, 60 mV, 80 mV, 100 mV, 120 mV. *I*_0_ represents the open current of single M113R/T115A nanopore. The black arrows indicate the addition of glycan. All measurements were performed in a symmetric buffer (3 M KCI, 10 mM Citric acid, pH 5.0), under an applied voltage at tra*ns* side. (*n*≥3)

**Figure S5.**
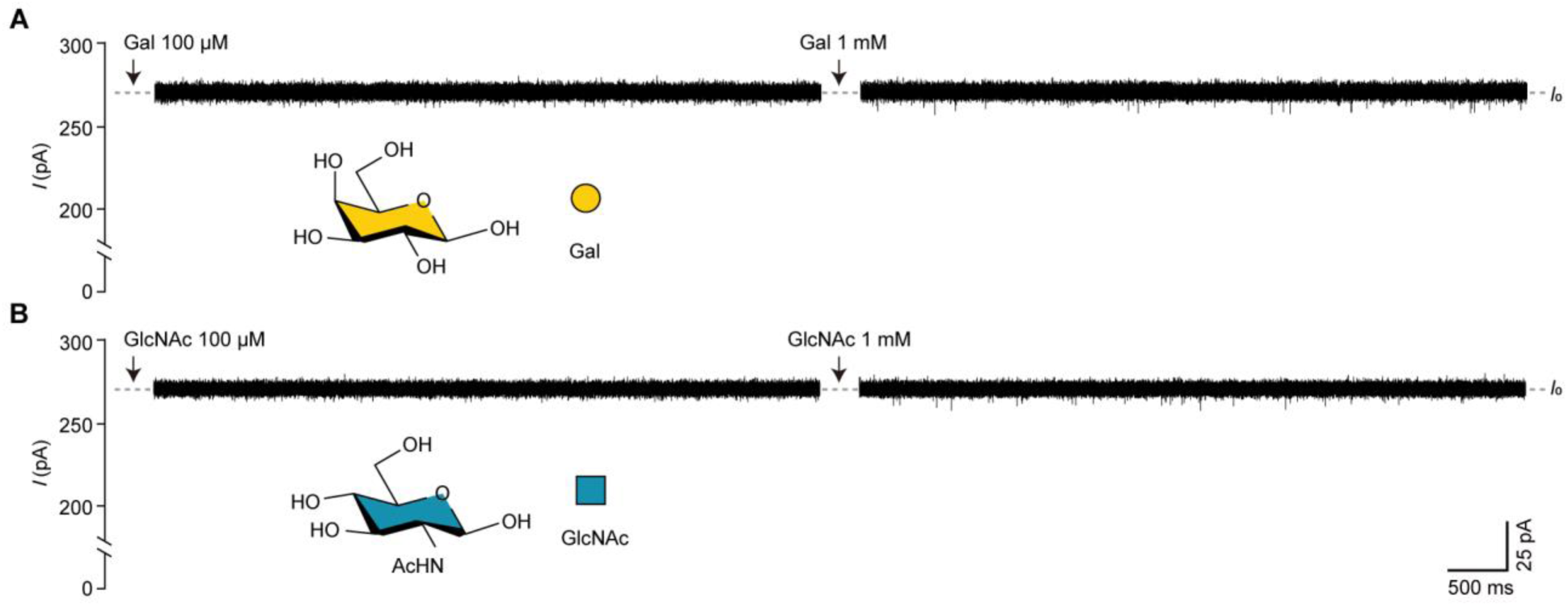
Detection of monosaccharides by M113R/T115A. (A) Representative ionic current traces of galactose (Gal) with concentration of 100 μM and 1 mM sensed by M113R/T115A. (A) Representative ionic current traces of N-Acetylglucosamine (GlcNAc) with concentration of 100 μM and 1 mM sensed by M113R/T115A. The monosaccharide was depicted with chemical structure and Symbol Nomenclature for Glycans (SNFG). The black arrows indicate the addition of glycan. All measurements were performed in a symmetric buffer (3 M KCI, 10 mM Citric acid, pH 5.0), under an applied +100 mV voltage at tra*ns* side. (*n*≥3)

**Figure S6.**
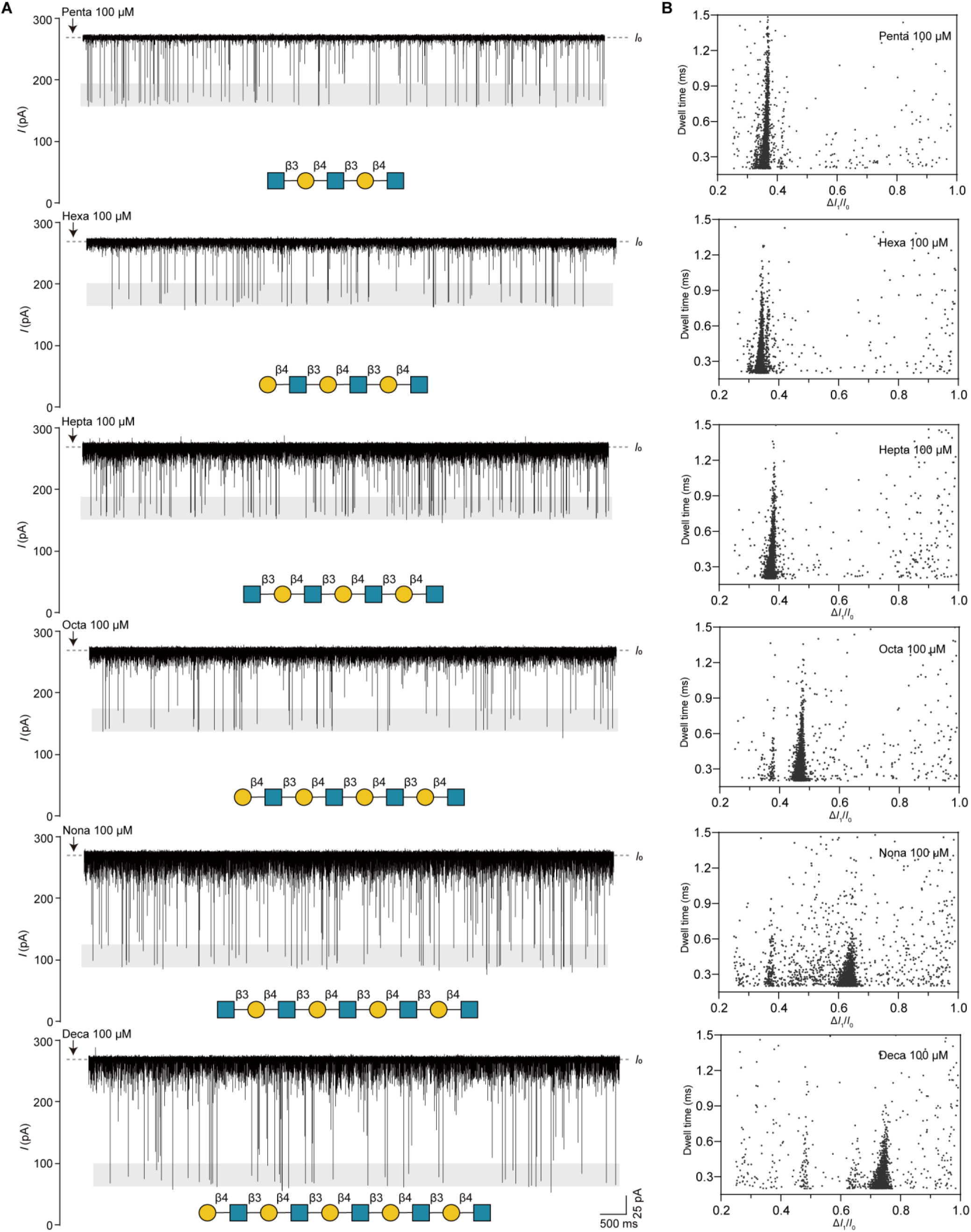
Detection of glycan standards from Pentasaccharide (Penta) to Decasaccharide (Deca) by M113R/T115A. (A) Representative ionic current traces after the addition of 100 μM glycan to *cis* side. Glycans were depicted with Symbol Nomenclature for Glycans (SNFG). The black arrows indicate the addition of glycan. (C) The scatter plots of Δ*I*_1_/*I*_0_ versus Dwell time of Penta, Hexa, Hepta, Nona, Deca. All measurements were performed in a symmetric buffer (3 M KCI, 10 mM Citric acid, pH 5.0), under an applied +100mV voltage at tra*ns* side. (*n*≥3)

**Figure S7.**
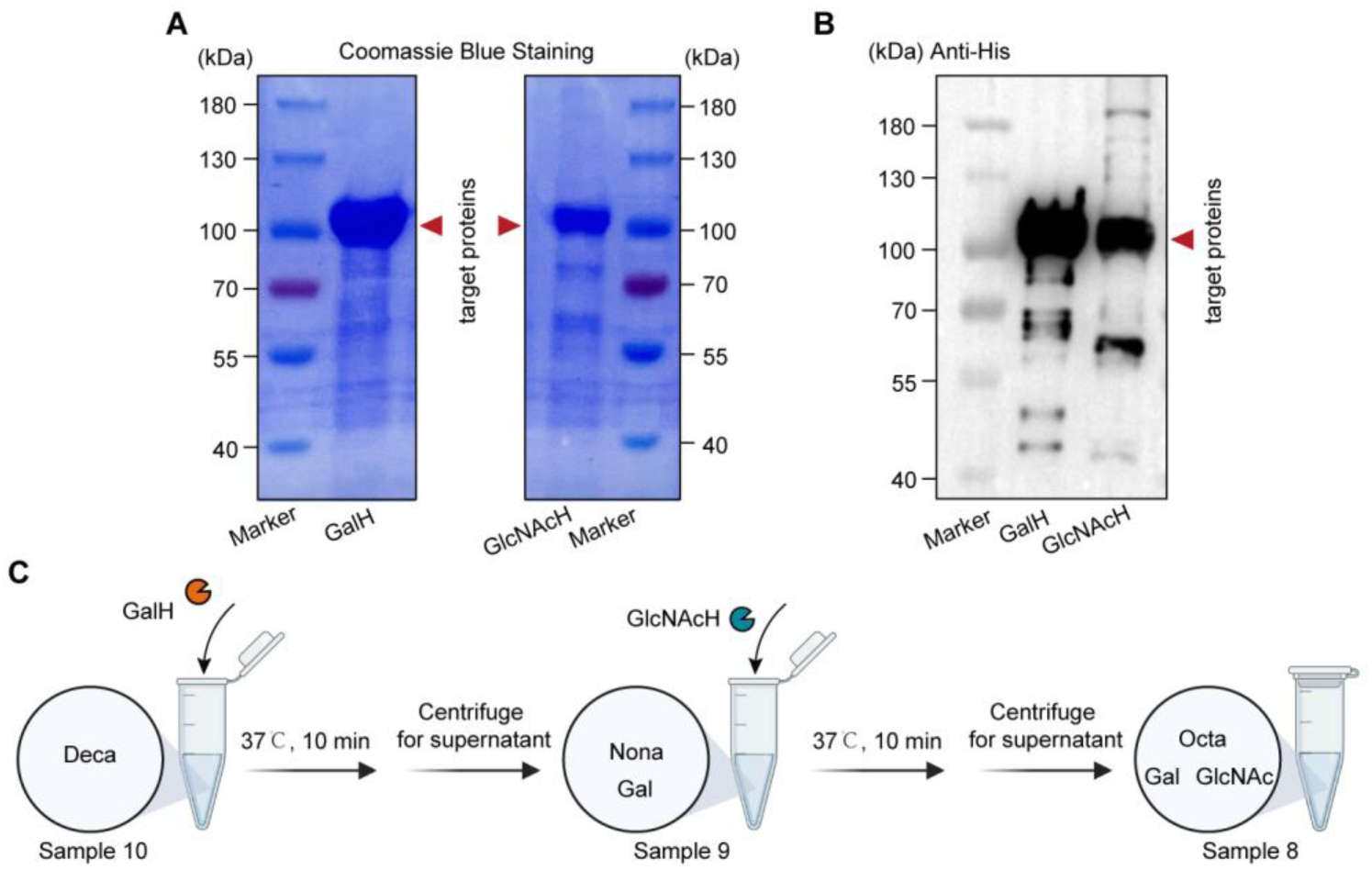
Purification of glycosidase protein of GalH and GlcNAcH. (A) 12% SDS-PAGE gel electrophoresis with coomassie blue staining. (B) Western blot probed with anti-his-tag antibody. (C) The scheme of hydrolysis experiment of decasaccharide using GlcNAcH and GalH. A 500 μL mixture containing 5 mM of substrate decasaccharide and 0.5 μg/μL GalH was incubated at 37°C for 10 min. Then, the reaction was quenched by heating in boiling water for 10 min. Insoluble impurities were removed by centrifugation (10000 rpm, 5 min). The supernatant was transferred to a tube as Sample 9 (S9). Afterwards, next exoglycosidase of GlcNAcH was added in S9. Sample 8 were obtained via similar procedure.

**Figure S8.**
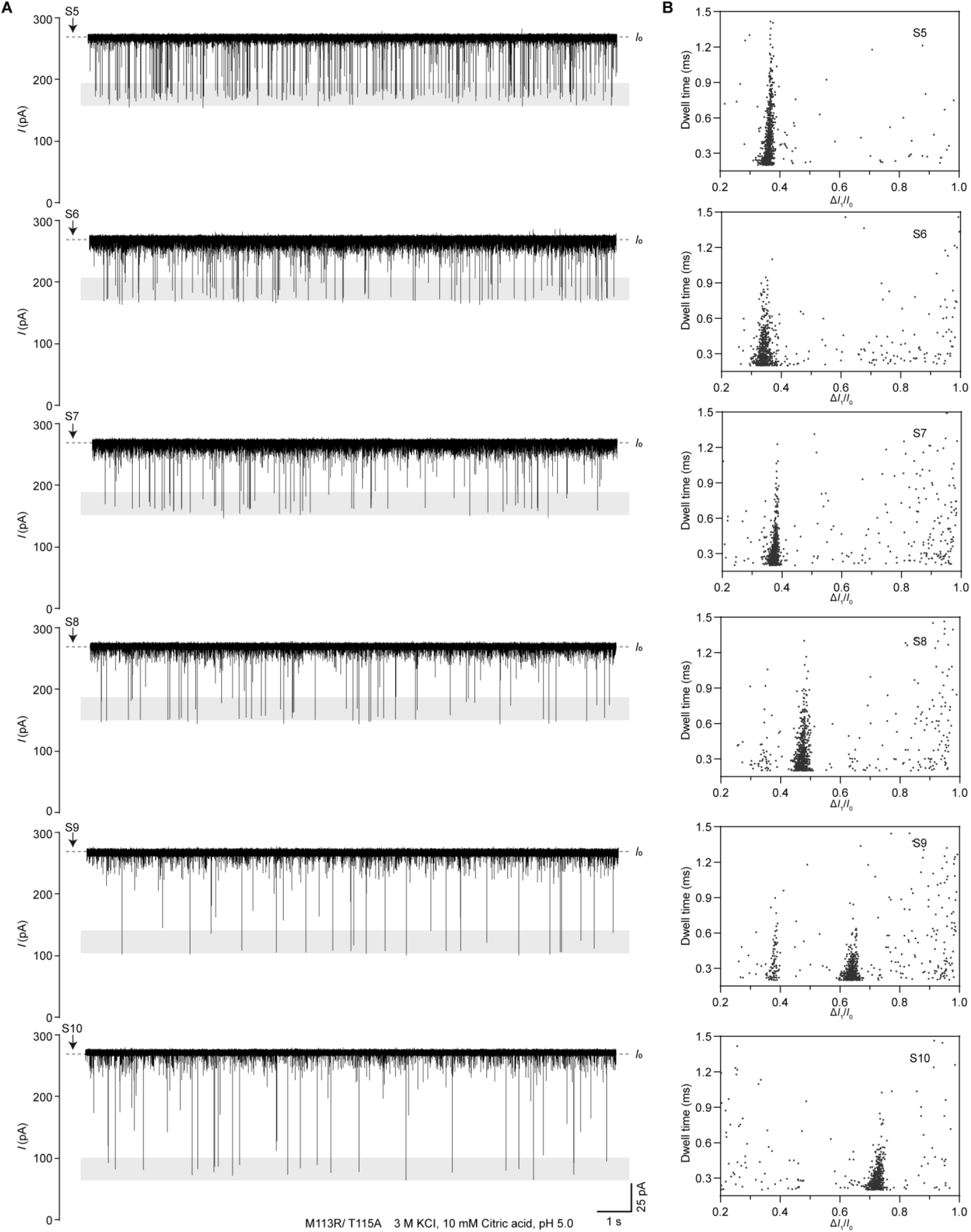
Detection of glycan hydrolysis samples. (A) Representative ionic current traces after the addition of 5.2 μL sample (S10, S9, S8, S7, S6, S5) to *cis* side (The total final glycan concentration is 100 μM). The black arrows indicate the addition of samples. (B) The scatter plots of Δ*I*_1_/*I*_0_ versus Dwell time of S10, S9, S8, S7, S6, S5. All measurements were performed in a symmetric buffer (3 M KCI, 10 mM Citric acid, pH 5.0), under an applied +100mV voltage at tra*ns* side. (*n*≥3)

**Figure S9.**
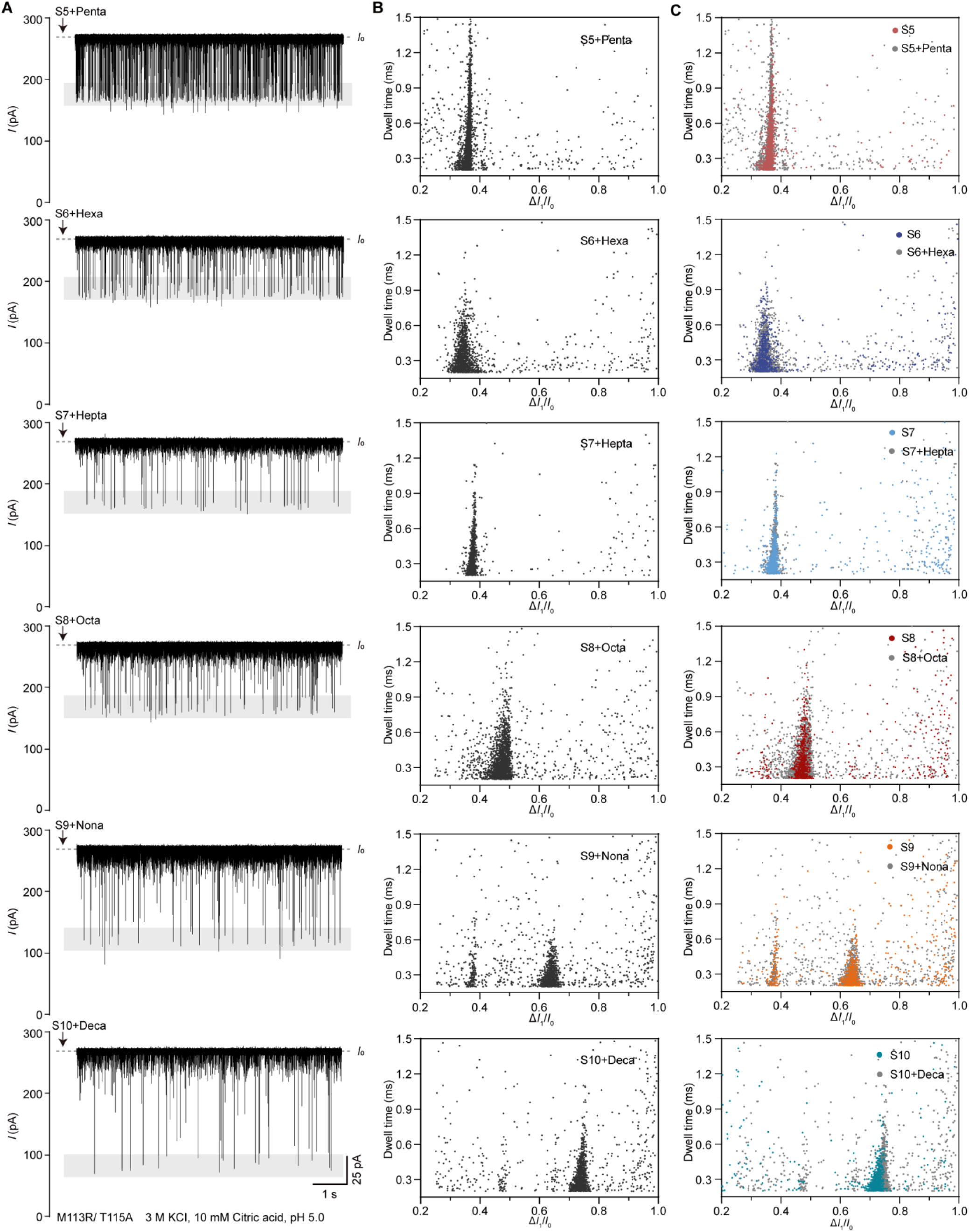
Detection of mixture of glycan hydrolysis samples and glycan standards of target products. (A) Representative ionic current traces after the addition of 5.2 μL sample (S10, S9, S8, S7, S6, S5) and glycan standards of target products (Penta, Hexa, Hepta, Octa, Nona, Deca, with final concentration of 100 μM) to *cis* side (The total final glycan concentration is 100 μM). The black arrows indicate the addition of samples. (B) The scatter plots of Δ*I*_1_/*I*_0_ versus Dwell time of S10+Deca, S9+Nona, S8+Octa, S7+Hepta, S6+Hexa, S5+Penta. (B) Merge of Δ*I*_1_/*I*_0_ versus Dwell time scatter plots of S10+Deca, S9+Nona, S8+Octa, S7+Hepta, S6+Hexa, S5+Penta with S10, S9, S8, S7, S6, S5, respectively. All measurements were performed in a symmetric buffer (3 M KCI, 10 mM Citric acid, pH 5.0), under an applied +100 mV voltage at *trans* side. (*n*≥3)

**Figure S10.**
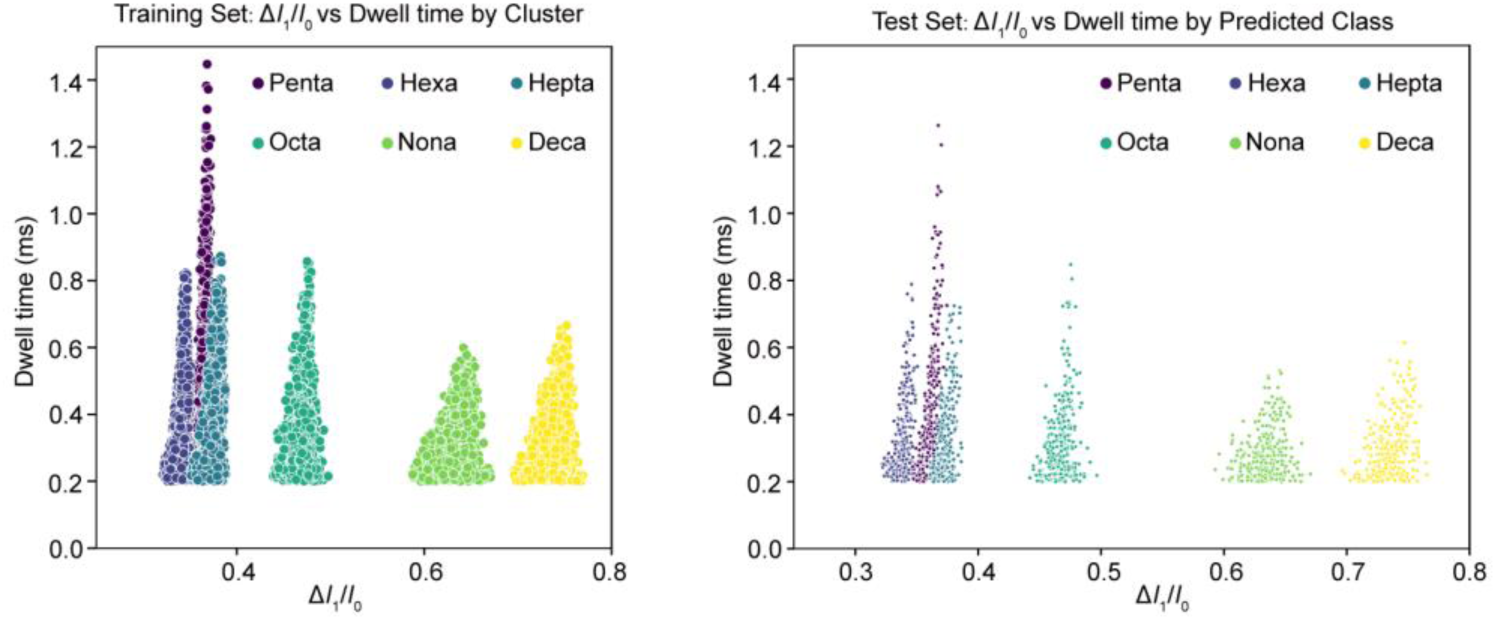
Data preparation for machine learning. (A) Training set: Δ*I*_1_/*I*_0_ versus Dwell time by cluster. (B) Test set: Δ*I*_1_/*I*_0_ versus Dwell time by predicted class. A minimum of 1500 events of each glycan standards were included after ignoration of 0.2 ms and filtration via Agglomerative clustering algorithm, with 80% used in the training set for model training and 20% used as the test set for model testing.

**Figure S11.**
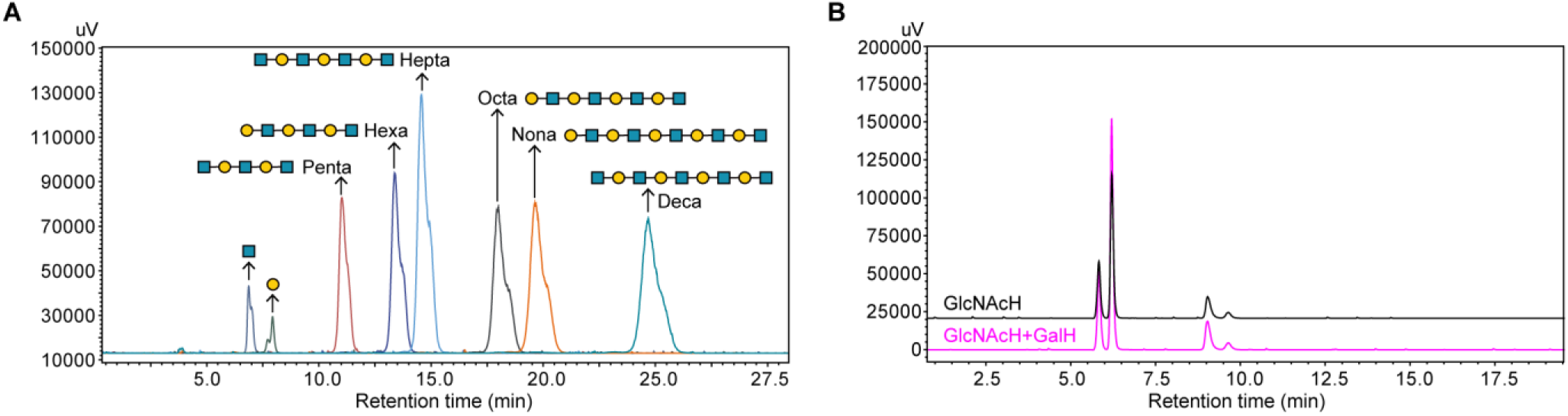
HPLC profile of the glycans and enzymes. (A) The concentration of each glycan is 5 mM. HPLC chromatogram showing significant retention time differences between the glycans. The retention time of GlcNAc, Gal, Penta, Hexa, Hepta, Octa, Nona, and Deca sequentially was 6.83±0.06 min, 7.73±0.06 min, 10.87±0.12 min, 13.13±0.23 min, 14.30±0.35 min, 17.67±0.47 min, 19.23±0.58 min, 24.03±0.58 min. HPLC profiles of the glycans displaying two partially overlapping peaks represent the α and β anomer. (B) HPLC profile of the enzyme solution. GlcNAcH, control of reaction sample containing 0.5 μg/μL GlcNAcH (without any glycan). GlcNAcH+GalH, control of reaction sample containing 0.5 μg/μL GlcNAcH and 0.5 μg/μL GalH (without any glycan).

**Figure S12.**
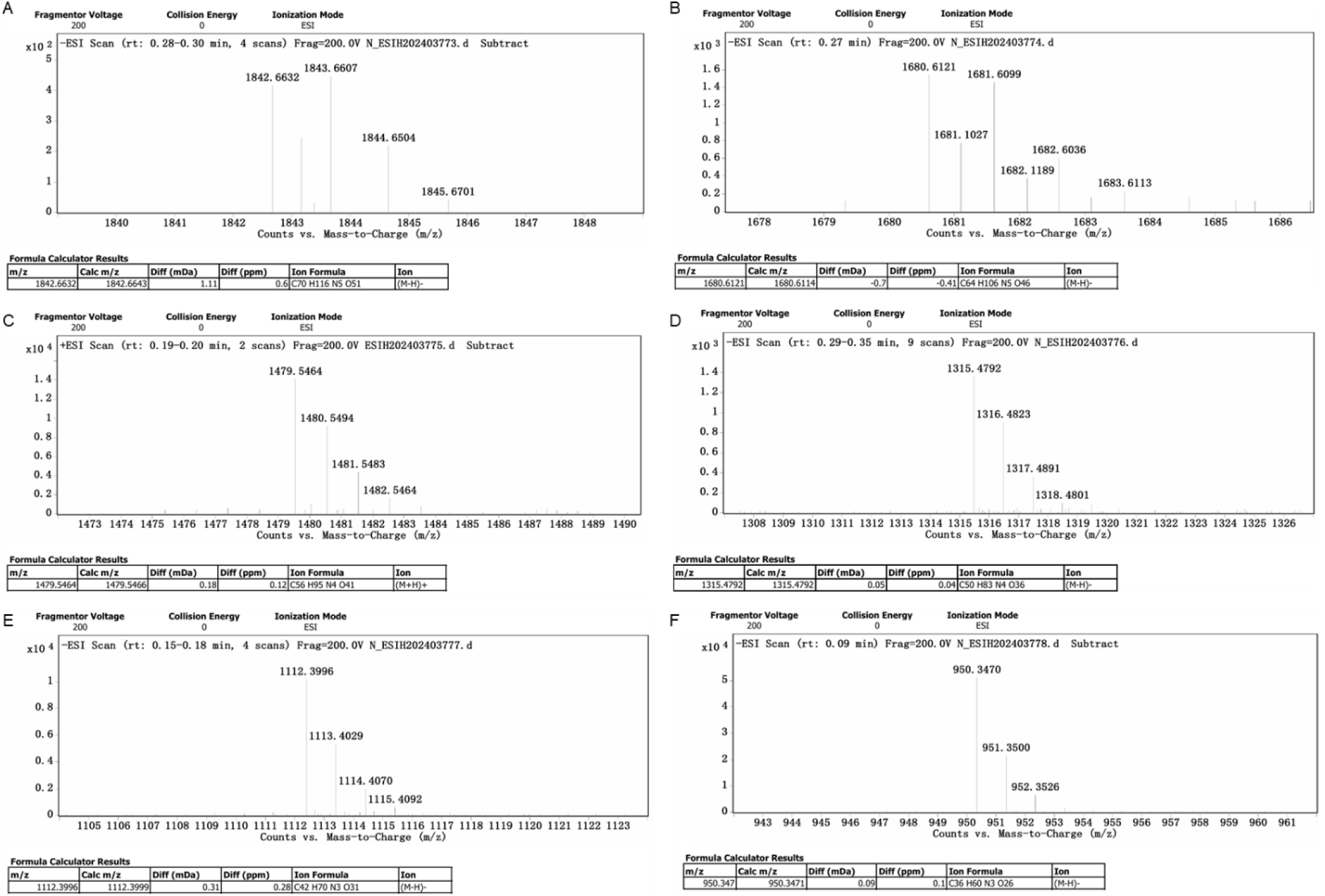
Identification of glycan hydrolysis sample via Mass Spectrum. (A) ESI-HR profile of Sample 10 (S10). The solution of Sample 1 was analyzed by ESI-HR and showed to decasaccharide (*m/z* calcd for C_70_H_116_N_5_O_51_ [M-H]^-^ 1842.6632, found 1842.6632). (B) ESI-HR profile of Sample 9 (S9). The solution of Sample 9 was analyzed by ESI-HR and showed to nonasaccharide (*m/z* calcd for C_64_H_106_N_5_O_46_ [M-H]^-^ 1680.6121, found 1680.6121). (C) ESI-HR profile of Sample 8 (S8). The solution of Sample 8 was analyzed by ESI-HR and showed to octasaccharide (*m/z* calcd for C_56_H_95_N_4_O_41_ [M+H]^+^ 1479.5464, found 1479.5464). (D) ESI-HR profile of Sample 7 (S7). The solution of Sample 7 was analyzed by ESI-HR and showed to heptasaccharide (*m/z* calcd for C_50_H_83_N_4_O_36_ [M-H]^-^ 1315.4792, found 1315.4792). (E) ESI-HR profile of Sample 6 (S6). The solution of Sample 6 was analyzed by ESI-HR and showed to hexasaccharide (*m/z* calcd for C_42_H_70_N_3_O_31_ [M-H]^-^ 1112.3996, found 1112.3996). (F) ESI-HR profile of Sample 5 (S5). The solution of Sample 5 was analyzed by ESI-HR and showed to Pentasaccharide (*m/z* calcd for C_36_H_60_N_3_O_26_ [M-H]^-^ 950.347, found 950.347).

## Supplementary Tables

**Table S1.**
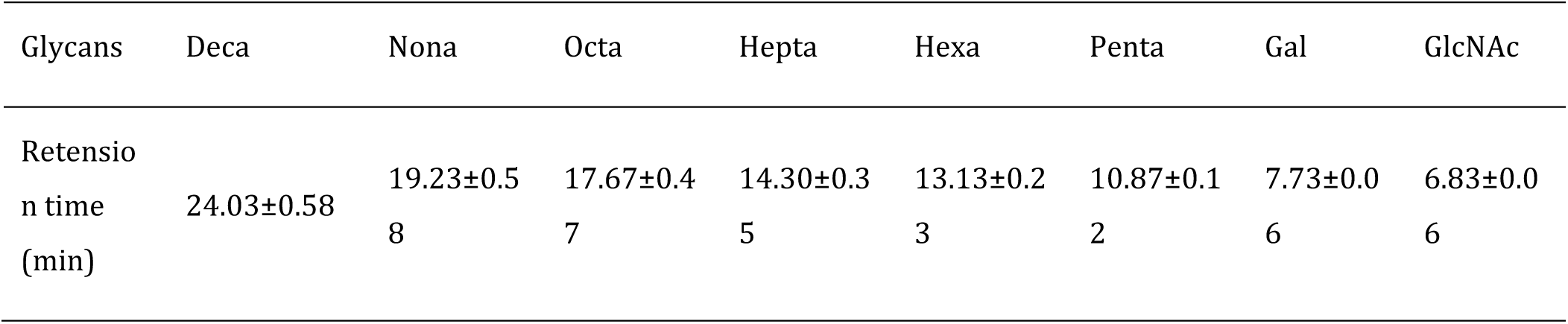
Statistics of retention time of glycan standards. (*n*≥3)

**Table S2.**
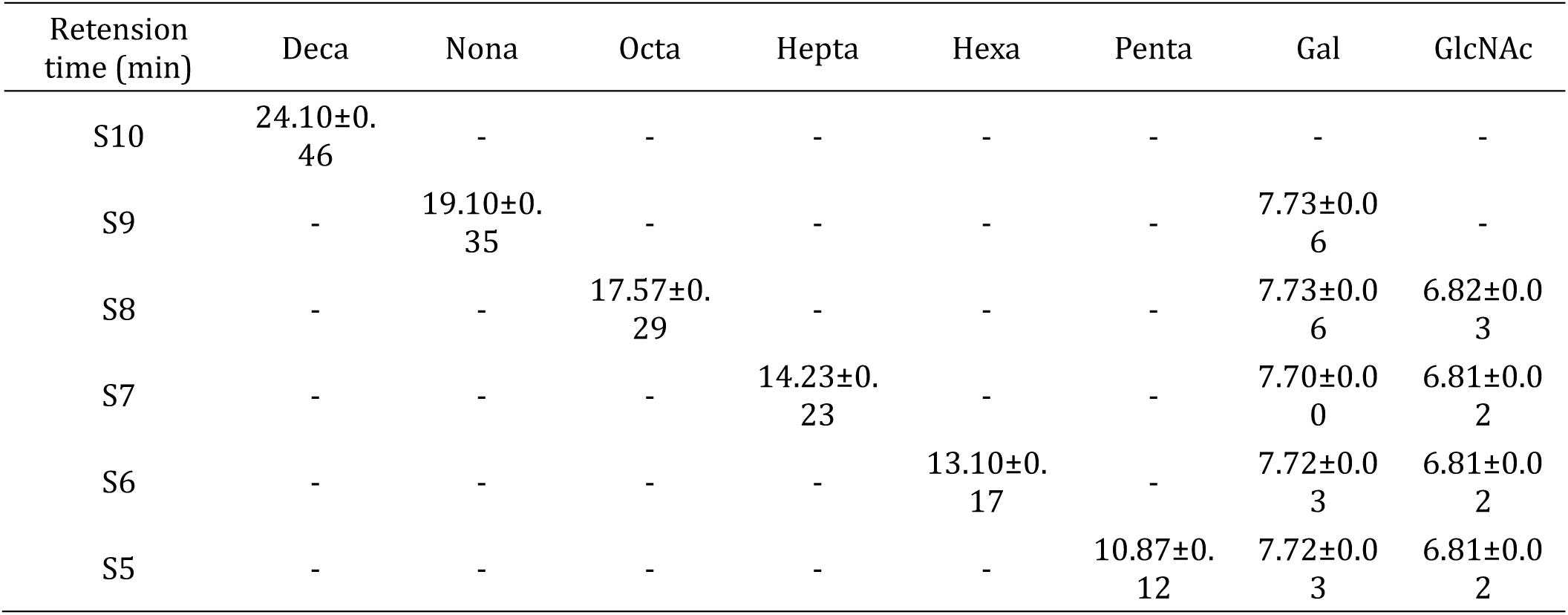
Statistics of retention time of glycan hydrolysis samples. (*n*≥3)

